# Three-dimensional connectivity and chromatin environment mediate the activation efficiency of mammalian DNA replication origins

**DOI:** 10.1101/644971

**Authors:** Karolina Jodkowska, Vera Pancaldi, Ricardo Almeida, Maria Rigau, Osvaldo Graña-Castro, José M. Fernández-Justel, Sara Rodríguez-Acebes, Miriam Rubio-Camarillo, Enrique Carrillo-de Santa Pau, David Pisano, Fátima Al-Shahrour, Alfonso Valencia, María Gómez, Juan Méndez

## Abstract

In mammalian cells, chromosomal replication starts at thousands of origins at which replisomes are assembled and bidirectional DNA synthesis is established. The slowdown of DNA polymerases at endogenous or exogenous obstacles triggers the activation of additional ‘dormant’ origins whose genomic positions and regulation are not well understood. Here we report a comparative study of origin activity in mouse embryonic stem cells growing in control conditions or in the presence of mild replication stress. While stress-responsive origins can be identified, we find that the majority of them are also active, albeit with lower frequency, in the control population. To gain insights into the molecular and structural determinants of origin efficiency, we have analyzed the genetic and epigenetic features of origins stratified according to their frequency of activation. We have also integrated the linear origin maps into three-dimensional (3D) chromatin interaction networks, revealing a hierarchical organization in which clusters of connected origins are brought together by longer-range chromatin contacts. Origin efficiency is proportional to the number of connections established with other origin-containing fragments. Interacting origins tend to be activated with similar efficiency and share their timing of replication even when located in different topologically associated domains. Our results are consistent with a model in which clusters of origins are arranged in 3D in replication factories. Within each factory, ‘main’ and ‘dormant’ origins are functionally defined by a combination of chromatin environment and 3D connectivity.

## INTRODUCTION

Proliferating cells duplicate their genetic information before each cell division. The copying of mammalian DNA chromosomes starts at thousands of replication origins that serve as assembly points for the protein machinery responsible for DNA synthesis. Origins are fundamental elements for genomic stability, and their precise number, position and regulation has been subject to decades of investigation (reviewed by Aladjem and Redon, 2016; Prioleau and MacAlpine, 2016; Rivera-Mulia and Gilbert, 2016). Chromosomal segments replicate in S phase following a temporal order which is commonly referred to as the “replication timing” (RT) program. While its biological significance is not fully understood, the RT program is evolutionary conserved and has been linked to origin activation, large-scale chromatin folding and nuclear compartmentalization (reviewed by Rhind and Gilbert, 2013).

Different technical approaches have been used to identify origins at the genome-wide level, including the isolation of ‘short nascent DNA strands’ (Cadoret et al, 2008; Sequeira-Mendes et al, 2009; Martin et al, 2011; Besnard et al, 2012; Picard et al, 2014; Cayrou et al, 2011, 2015; Almeida et al, 2018); the capture and sequencing of origin-containing replication ‘bubbles’ (Mesner et al, 2013); the analysis of the strand distribution of Okazaki fragments (McGuffee et al, 2013; Petryk et al, 2016; Chen et al, 2019); sequencing of newly-synthesized labeled DNA (Langley et al, 2016; Macheret and Halazonetis, 2018; Tubbs et al, 2018); and chromatin immunoprecipitation of origin-binding proteins (Dellino et al, 2013; Miotto et al, 2016; Sugimoto et al, 2018). Depending on the resolution obtained, these methods have identified putative individual origins or broader initiation zones in several cell lines. While a consensus map of individual origin positions is yet to be reached, common patterns have emerged from these studies, such as the frequent localization of origins at transcription start sites (TSS) and CpG islands (CGIs). G-quadruplex structures have also been identified in the vicinity of origins (Besnard et al, 2012; Cayrou et al, 2011, 2015; Picard et al, 2014; Comoglio et al, 2015) and may contribute to origin function (Valton et al, 2014). A recent analysis of single-nucleotide polymorphisms has also identified a short (~40 bp) region around origins with a deficiency in common variants and indels, indicative of strong selective pressure. This signature is observed at all origins, including those not containing CGI, TSS, or G4 structures (Massip et al, 2019).

DNA replication displays flexibility in different systems, including Drosophila and mammalian cells (Besnard et al, 2012; Picard et al, 2014; Cayrou et al., 2015; Comoglio et al, 2015). Besides the fact that different cell types may use origins with different efficiencies, pioneering work by J.H. Taylor (1977) described how cells artificially held in S phase increased the number of DNA replication sites due to the activation of new origins. Years later, this concept could be integrated with the fact that more origins are ‘licensed’ by initiator proteins ORC, CDC6, CDT1 and MCM2-7 than those actually needed to duplicate the genome (reviewed by Alver et al, 2014; Shima and Pederson, 2017). Thus, many origins remain in a dormant state and are passively replicated by active forks in S phase. However, dormant origins can be activated in situations of ‘replicative stress’ (RS), i.e. when forks are slowed or stalled by DNA lesions, conflicts with the transcriptional machinery or other factors (reviewed by Hamperl and Cimprich, 2016; Muñoz and Mendez, 2017). Stress-responsive origins provide a compensatory mechanism to complete duplication in mammalian cells (Ge et al, 2007; Ibarra et al, 2008) and their relevance *in vivo* has been demonstrated in mouse strains with reduced expression of origin licensing proteins MCM2-7, which are viable but suffer from stem cell deficiencies, anemia and cancer (Shima et al, 2007; Pruitt et al, 2007; Kawabata et al, 2011; Alvarez et al, 2015). A full complement of MCM proteins is also needed to maintain the functionality of hematopoietic stem cells (Flach et al, 2014). Of note, the availability of extra origins may also pose a risk upon certain oncogenic stimuli that induce promiscuous origin activity, resulting in DNA breaks caused by a higher frequency of collisions between replication and transcription (Macheret and Halazonetis, 2018). A better understanding of these processes requires in-depth information about the genomic positions and regulation of common and stress-responsive origins.

In this study, we have mapped origin activity in embryonic stem cells (ESCs) in the absence or presence of mild RS to trigger dormant origin activation. The majority of responsive origins correspond to normal origins that were activated with low efficiency in the population. Through extensive cross-analyses of high- and low-efficiency origins with genetic and epigenetic features, as well as the integration of origins in three-dimensional chromatin interaction networks, we propose new determinants of origin efficiency that can functionally separate main from dormant origins.

## RESULTS

### Mapping mESC replication origins under stress

To identify active origins in mouse embryonic stem cells (mESCs) with high resolution, we used deep sequencing of <u>s</u>hort <u>n</u>ascent <u>s</u>trands (SNS-Seq; **Supp Figure 1A**), a method that has yielded reproducible results in different laboratories. SNS-Seq was conducted in normal growth conditions (hereafter referred to as “WT”) or in two experimental settings that trigger the activation of extra origins: (i) the presence of DNA polymerase inhibitor aphidicolin (APH); (ii) ectopic expression of CDC6, a limiting factor for origin licensing and activation (Muñoz et al, 2017). Two experimental replicates of each condition were analyzed. The treatment with APH mimics RS, i.e. it slows down replication forks and triggers the activation of extra origins as a compensatory mechanism. In turn, CDC6 overexpression aims at enhancing origin activity directly, with fork slowdown being a likely consequence of reduced dNTP availability (Rodriguez-Acebes et al, 2018). In the experimental conditions used, both APH treatment and CDC6 overexpression slowed down forks without blocking overall DNA synthesis or inducing significant DNA damage (**Supp Figure 1B**). The quality of SNS preparations was monitored by controlling the completeness of lambda-exonuclease digestion, a necessary step to eliminate contaminant DNA (Foulk et al, 2015; **Supp Figure 1C**), as well as confirming the enrichment of SNS at a known origin relative to its flanking region (Sequeira-Mendes et al, 2009; **Supp Figure 1D**).

Two separate algorithms were used to identify peaks from SNS-Seq aligned reads: MACS, a ChIP-Seq tool that has been applied to map origins from SNS-Seq data (Comoglio et al, 2015), and a dedicated algorithm optimized for SNS-Seq by Picard *et al* (2014) that takes into account local coverage heterogeneities (**Figure 1A**). Overall, the total number of peaks was higher when called by MACS (range 71,435-94,747 compared with 41,376-94,893 with the Picard algorithm; **Table 1**), reflecting the higher stringency of the latter. In one of the samples (WT-I), in which the signal-to-noise ratio was particularly high, both algorithms identified a similar number of peaks (**Table 1**). Reproducibility between biological replicates was assessed by pairwise correlation of origin number per genomic segment, and by the distribution of SNS-seq reads for both replicates around the peak centers of one of them (**Figure 1B)**. Peak overlap between replicates was in the 76-82% range, in line to what has been reported in other studies, except between the two WT samples (49-54% depending on the algorithm), likely reflecting an unusually high signal-to-noise ratio in one of the replicates. To minimize the influence of technical variability in subsequent analyses, only those peaks called by both algorithms in the two replicates were included in the origin datasets. With these stringent criteria, 20,174, 31,685 and 31,402 active origins were defined in WT, APH and CDC6 conditions, respectively (**Supp Table 1**).

**Figure 1.**
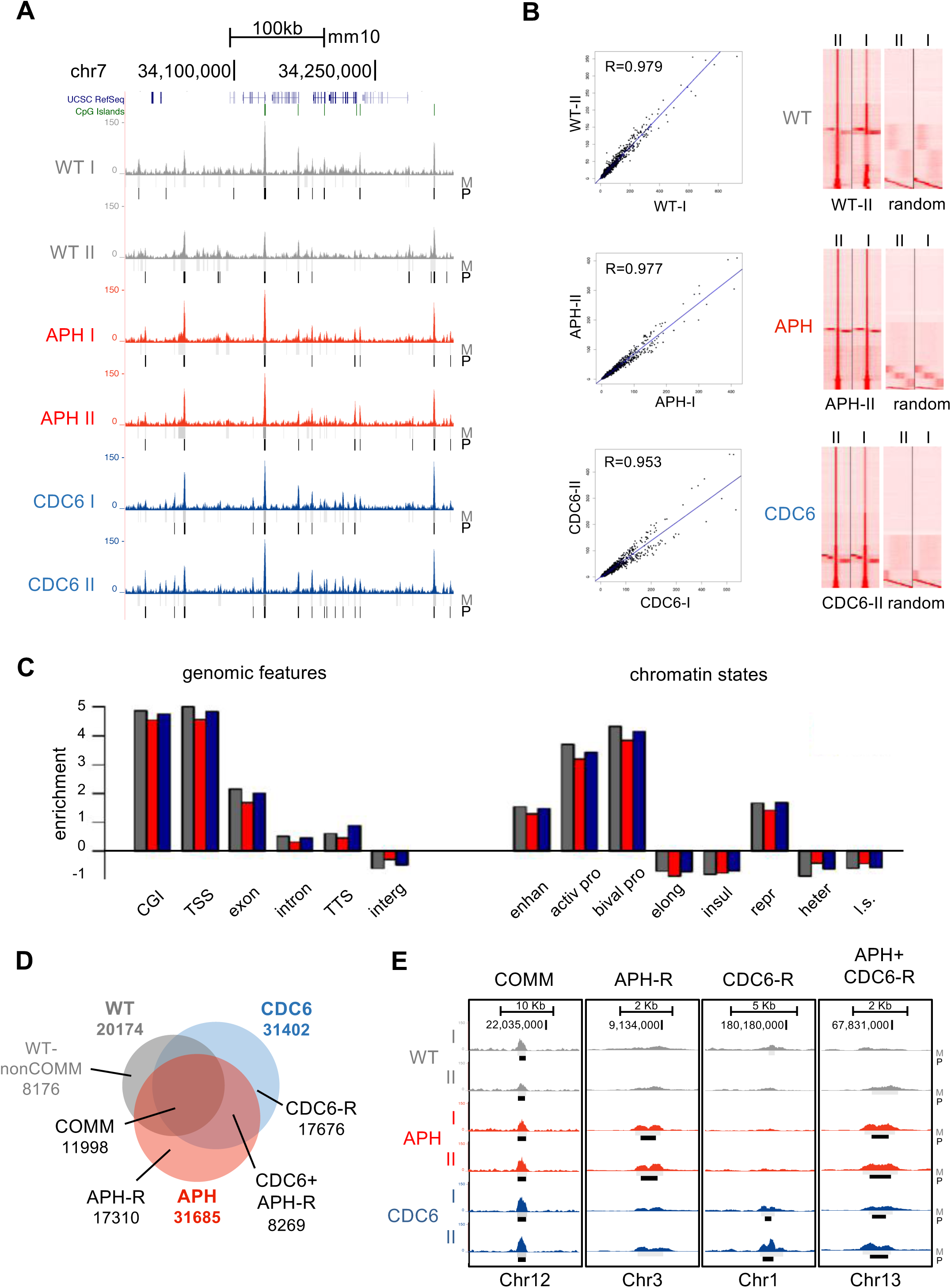
Genome-wide mapping and features of mESC replication origins in normal and stress conditions. **A.** Genome browser image showing read density tracks in a representative fragment of chromosome 7 from two SNS-Seq replicates in control mESCs (WT, grey), mESCs treated with aphidicolin (APH, red) and mESCs after CDC6 overexpression (CDC6, blue). Vertical dashes indicate the positions of peaks called by MACS (M, grey) and Picard (P, black) algorithms. **B.** Left, pairwise correlation of origin number per genomic segment between replicates. Right, heatmap distribution of WT, APH or CDC6 SNS-seq reads from experimental replicates I and II around the origin peak centres in one of them (II) as defined by the Picard algorithm. Read distribution is also shown around equivalent sets of randomised origin peak centres. **C.** Enrichment of WT (grey), APH (red) and CDC6 (blue) origins at the indicated genomic features (left) or chromatin states (right), relative to the randomized controls. CGI, CpG island; TSS, transcription start site; TTS, transcription termination site; interg, intergenic region; enhan, enhancer; activ pro, active promoter; bival pro, bivalent promoter; heter, heterochromatin; elong, transcriptional elongation; insul, insulator; repr, repressed chromatin; l.s., low signal. All enrichments are significant at p<0.001. **D.** Venn diagram of origin subsets, determined by intersections of WT, APH and CDC6 datasets. **E.** Genome browser examples of common and stress-responsive origins.

**Table 1.** Origin identification and genomic coverage using MACS and Picard algorithms.

|  | **aligned**<br>**SNS reads,**<br>**nr** | **origin**<br>**nr,**<br>**MACS** | **mean**<br>**length**<br>**(Kb)** | **cumul**<br>**length**<br>**(Mb)** | **genome**<br>**covered**<br>**%** | **origin**<br>**nr,**<br>**PICARD** | **mean**<br>**length**<br>**(Kb)** | **cumul**<br>**length**<br>**(Mb)** | **genome**<br>**covered**<br>**%** | **shared**<br>**origins,**<br>**nr** | **shared**<br>**origins,**<br>**%** |
| --- | --- | --- | --- | --- | --- | --- | --- | --- | --- | --- | --- |
| **WT-I** | 57,789,135 | 94,747 | 1.60 | 152 | 5.77 | **94,893** | **0.667** | **63.3** | **2.40** | 65,623 | 69.1 |
| **WT-II** | 64,910,836 | 81,056 | 1.67 | 135 | 5.12 | **44,860** | **0.818** | **36.7** | **1.39** | 41,974 | 93.5 |
| **APH-I** | 33,994,540 | 74,540 | 2.10 | 156 | 5.94 | **49,061** | **0.735** | **36.1** | **1.37** | 42,536 | 86.6 |
| **APH-II** | 42,562,298 | 71,435 | 1.81 | 129 | 4.91 | **41,376** | **0.770** | **31.8** | **1.21** | 38,200 | 92.2 |
| **CDC6-I** | 77,850,848 | 72,384 | 1.54 | 111 | 4.22 | **42,791** | **0.821** | **35.1** | **1.33** | 39,883 | 93.1 |
| **CDC6-II** | 60,639,981 | 82,071 | 1.39 | 114 | 4.33 | **62,922** | **0.718** | **45.2** | **1.71** | 57,020 | 90.5 |

### Origin enrichment at genomic elements and chromatin states

WT, APH and CDC6 origin datasets overlapped with CGIs, TSS and exons with much higher frequency than expected by chance (**Figure 1C, left**). Intersection analyses with different ‘chromatin states’ defined by epigenetic features in ESCs (Ernst and Kellis, 2010; Filion et al, 2010, Juan et al, 2016) confirmed that origins from all groups correlated preferentially with enhancers, promoters (active and bivalent) and also with the Polycomb repressed state. In contrast, insulators, heterochromatin and the ‘transcriptional elongation’ state contained origins with lower frequency than expected by chance (**Figure 1C, right**). Origins strongly correlated with histone modifying enzymes KDM2A-B, HDAC1-2, and MLL; histone marks H3K9ac, H3K4me2 and me3; components of the mediator and cohesin complexes; the TET1 DNA demethylase complex; components of transcriptional and chromatin remodeling complexes Polycomb, CoRest, NuRD and Sin3; RNA polymerase II and its promoter-located variants RNAPII-S5 and RNAPII-S7; as well as several transcription factors including MYC, MAX, KLF4, E2F1 and OCT4 (**Supp Figure 2A**). This is one of the most comprehensive characterisation of individual marks linked to mammalian replication origins generated so far, and it emphasizes their association with a combination of genetic and epigenetic elements that regulate chromatin accessibility and gene expression.

### Responsive origins are active in control conditions

The intersection of WT, APH and CDC6 datasets (**Figure 1D**) revealed a large group of origins active in every condition, which we hence termed “common” (COMM). Additionally, subsets of origins were apparently responsive to aphidicolin (APH-R), CDC6 (CDC6-R) or both stimuli (APH+CDC6-R; **Figure 1E**), while a smaller subset of WT origins were not identified upon stress (WT-nonCOMM; **Figure 1D** **and Supp Table 1**). The identification of APH-R and CDC6-R origins at new positions seems consistent with a deterministic model of DNA replication, in which every cell makes use of a defined set of origins along S phase, while extra origins activated by stress are located at new genomic positions (**Figure 2A, left**). However, the majority of genomic positions corresponding to responsive origins also displayed a moderate enrichment of SNS-seq reads in WT cells and many of them were identified as origins by one of the two peak-calling algorithms in at least one of the replicates (**Figure 1E**; see **Supp Figure 2B** for more examples). Indeed, origin activity in WT cells at responsive positions was confirmed by heatmap representations of WT SNS-Seq reads around APH-R or CDC6-R peak centers (**Figure 2B**). These finding suggests the alternative possibility that dormant origins are actually active in a fraction of the cells in the unchallenged S phase. Upon stress, they are activated with higher efficiency, i.e. in a higher percentage of the population. This model of origin usage has been termed ‘stochastic’ to indicate that each cell may use a slightly different subset of all possible origins, and the frequency of activation of each individual origin is determined by rules of probability (**Figure 2A, right;** Bechhoefer and Rhind, 2012).

**Figure 2.**
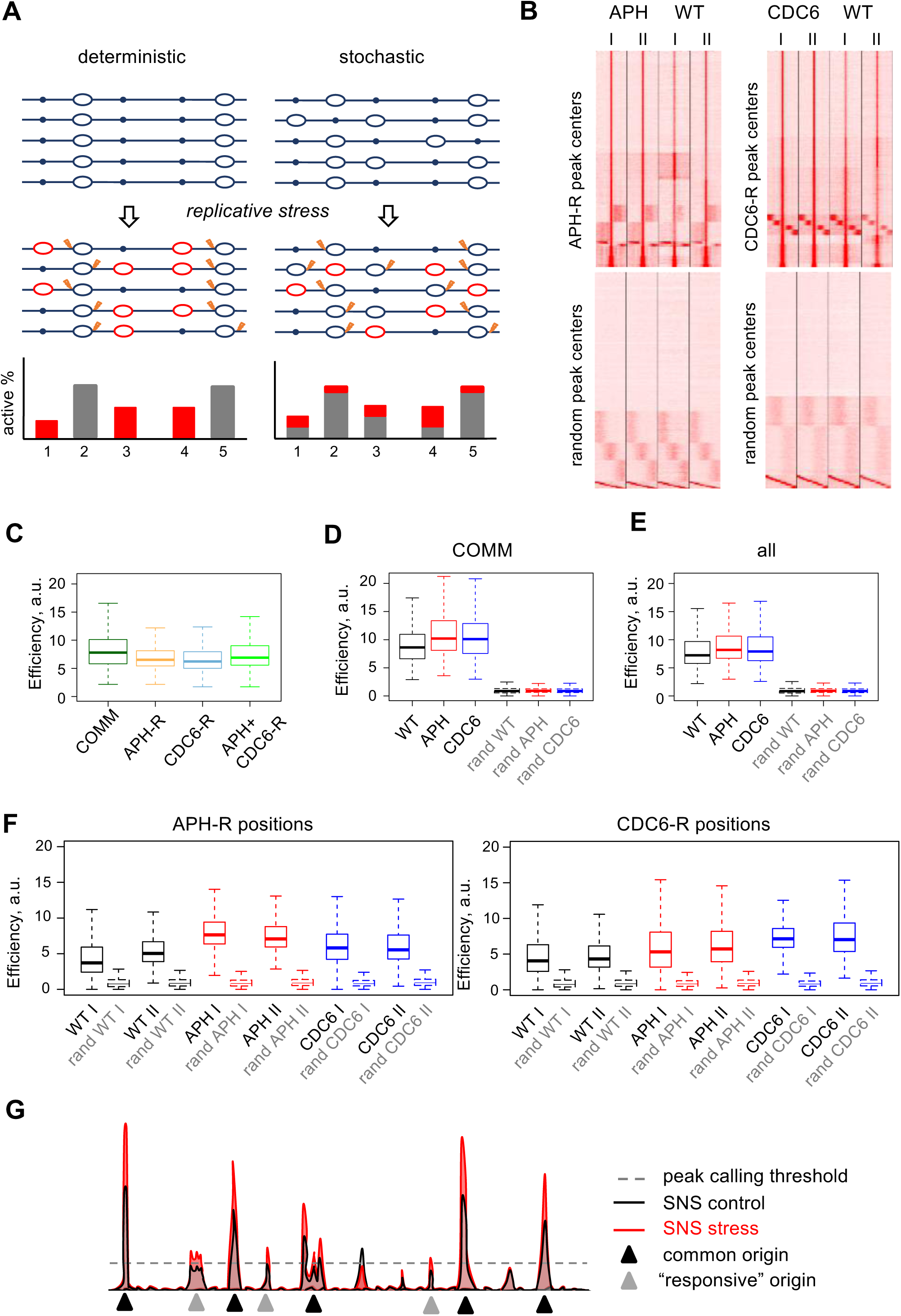
Increased activation of pre-existing origins upon stress. **A.** Schematic of deterministic vs stochastic models of origin use. A chromosomal region with five theoretical origins is represented. Bar graphs reflect the efficiency of the origins depicted above, in control conditions (grey) or after stress (red). **B.** Heatmap representation of the distribution of SNS-seq reads for the indicated experimental replicates around APH-R, CDC6-R or randomised peak centers (bottom panels). **C.** Box plots show the efficiencies of common and stress-responsive origins. Comparisons between all possible pairs of conditions are significant at p<10^−16^ (Wilcoxon signed-rank test). **D**. Efficiencies of common origins in WT, APH and CDC6 datasets. The efficiencies of randomized equivalent sets are also shown (dashed boxes). **E.** Same as (D) for the entire WT, APH and CDC6 origin datasets, or randomized equivalent sets. In the box-plots shown in C, D, E, median values are indicated by horizontal bars. **F.** Origin efficiency determined experimentally in each replicate of WT, APH or CDC6 conditions at the positions of APH-R (left) and CDC6-R (right) origins. The efficiency of equivalent sets of randomized origins is shown in dashed boxes. **G.** Schematic of efficiency profiles in control (black) and stress (red) conditions. The positions of common and responsive origins are based on the peak detection threshold (dashed line).

### Origin efficiency is increased upon mild replication stress

The frequency of activation of each origin in the population, termed origin efficiency, was estimated from the SNS read density of its corresponding peak in the different conditions used. In this type of analysis, a similar number of SNS-Seq reads from all conditions was used, and the average background signal in each track was taken into account (Methods). The efficiency of COMM origins (calculated for each origin as the average of its efficiency in the three experimental conditions) was higher than that of responsive origins of every category (**Figure 2C**), reflecting that they are active in a higher percentage of the cell population. The median efficiency value of COMM origins (calculated separately in WT, APH and CDC6 conditions) increased upon exposure to APH or CDC6, relative to WT cells (**Figure 2D**). This effect was also observed when the analysis was extended to all origins in the datasets (**Figure 2E**). Of note, no differences in efficiency were observed in equivalent sets of randomized genomic positions mimicking origins (**Figure 2D-E**). As anticipated from the genome browser images, origins located at APH-R or CDC6-R positions displayed lower efficiency in WT cells than upon APH or CDC6 stimuli (**Figure 2F**). We conclude that the APH-R, CDC6-R and APH+CDC6-R origin subsets represent low-efficiency initiation sites whose activity remained below the stringent threshold set by the combination of MACS and Picard algorithms in WT cells, but not upon stress (**Figure 2G**).

### Origin efficiency correlates with TSS proximity

In a stochastic model of origin activation, one of the key questions becomes the nature of the molecular determinants that regulate origin efficiency in the population. To gain insights into this issue, origins in the three datasets (WT, APH and CDC6) were subdivided into four quartiles according to their relative efficiencies. Origin strength directly correlated with their presence at CGIs, TSS, exons, enhancers, active and poised promoters (**Figure 3A-C**). In addition, stronger origins displayed a more marked association with individual epigenetic marks (**Supp Figure 2C**). In every dataset, origin efficiency was inversely proportional to their distance to the nearest TSS, i.e. most efficient origins were located very close to TSS; **Figure 3D-F**), in agreement with the frequent coupling between replication and transcription initiation (Sequeira-Mendes et al, 2009; Lombraña et al, 2013; Chen et al, 2019).

**Figure 3.**
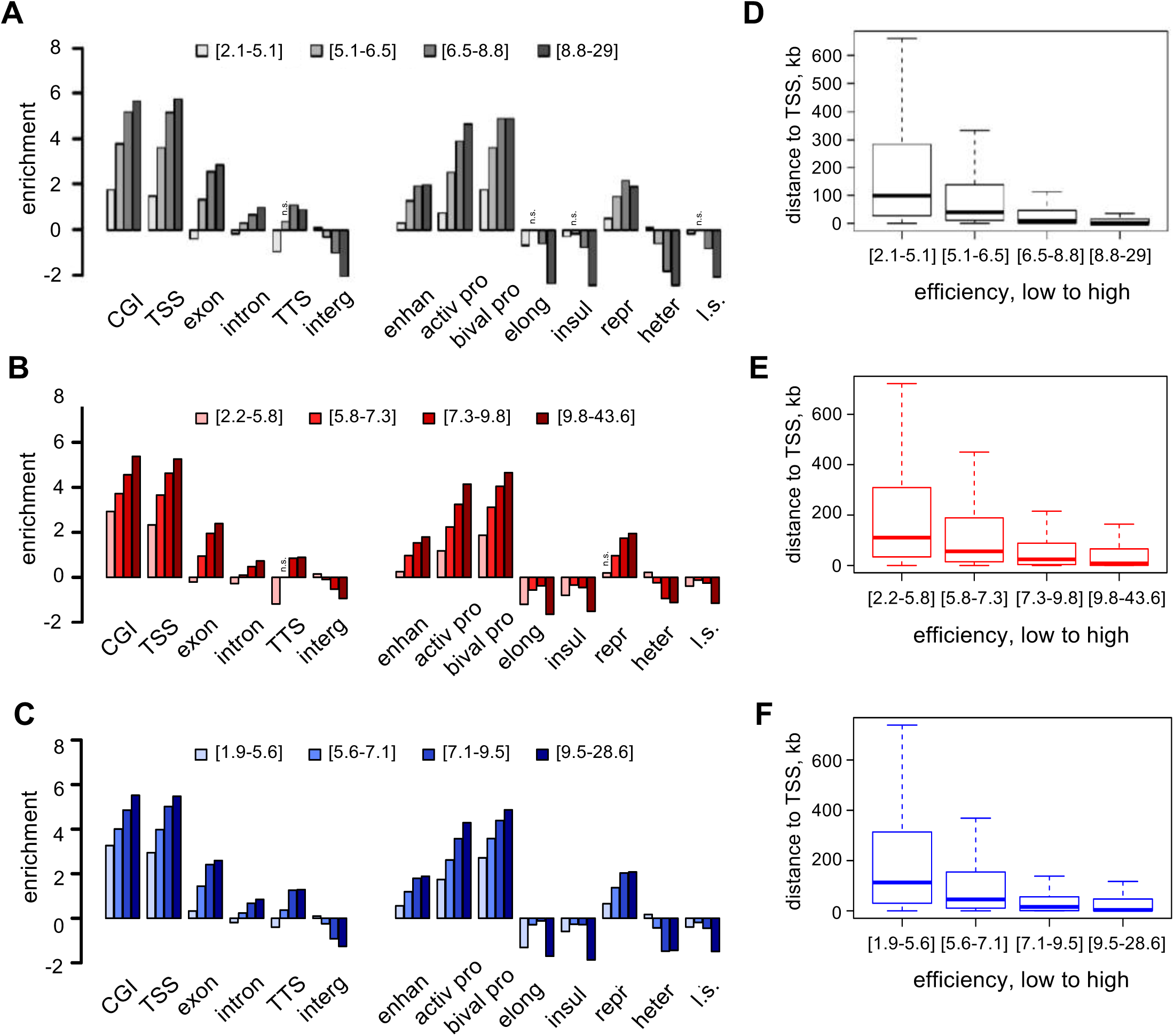
Origin efficiency correlates with genomic features and distance to TSS. **A-C.** Enrichment of WT (**A**), APH (**B**) or CDC6 (**C**) origins, distributed in quartiles according to their efficiencies, with genomic features (left) and chromatin states (right), relative to randomized controls. Labels as in **Figure 1**. All enrichments are significant at p<0.05 except for the ones marked with n.s. (not significant). **D-F.** Box plots showing the distribution of distances to nearest TSS of efficiency-stratified WT (**D**), APH (**E**) or CDC6 (**F**) origin datasets.

### Integration of origins into 3D chromatin contact maps

To provide a three-dimensional context for the replication start sites identified in this study, origin datasets were integrated with chromatin contact maps available for mESCs. Chromatin contact maps can be represented as networks in which chromatin fragments are located at the ‘nodes’ and experimentally-determined interactions between them are represented as ‘edges’ (Sandhu et al, 2012; Norton et al, 2018). Given the enrichment of origins at promoters, we focused our analysis on a Promoter-Capture HiC (PCHiC) map that identifies long-range contacts (>10 Kb, mean >1 Mb) between two promoters (P-P) or a promoter and a non-promoter region (P-O, “other end”; Schoenfelder et al, 2015). A large chromatin contact network can be derived from PCHiC data, involving >55,000 fragment nodes (mean length 5 Kb, range 0.15-60 Kb) and nearly 70,000 connections between them (edges; Pancaldi et al, 2016). Approximately 32% of WT origins overlapped with P fragments and 8% with O fragments. The vast majority of them (>95%) overlapped with a single chromatin fragment, and conversely, the majority of origin-containing PCHiC fragments hosted a single origin. Overall, up to 40% of WT origins localized at PCHiC nodes, which are by definition involved in 3D contacts between promoters, or between promoters and other ends (**Supp Table 1**). As expected, sets of randomized genomic positions mimicking WT origins overlapped with PCHiC nodes at a much lower frequency (13%). Given the frequent localization of origins at promoters, we devised and tested an alternative randomization method in which the distances of ‘random origins’ to their nearest TSS was arranged to be the same as for experimental origins. These sets overlapped with the network with a slightly lower frequency than experimental origins (33-34% after 20 randomizations, **Supp Table 2**), suggesting that the enrichment of origins at PCHiC chromatin fragments is mostly, but not exclusively, due to their overlap with promoter elements. A visual representation of WT origins in the PCHiC network can be explored in **Supp Figure 3A**.

### Origin presence is assortative in the P-P subnetwork

The PCHiC network can be divided into two subnetworks to separate contacts between promoters (P-P: 14,441 nodes; 20,515 edges) from contacts between promoters and other ends (P-O: 52,665 nodes; 49,472 edges). The distribution of WT origins in the P-P subnetwork is shown in **Figure 4A**. While this schematic representation does not indicate the actual positions of origins within the nucleus, it accurately represents the network of physical interactions between them (see inset in **Figure 4A**). This information enables the use of specific network analysis tools such as chromatin assortativity (ChAs; Pancaldi et al, 2016). ChAs is a correlation coefficient, ranging between −1 and 1, that measures the extent by which a given feature of any chromatin fragment is shared by the fragments that interact with it (**Figure 4B**). For instance, Polycomb Group (PcG) proteins, master regulators of mESCs genome architecture (Schoenfelder et al, 2015), and their associated histone marks display assortativity values ranging 0.2-0.35 in the PCHiC network (Pancaldi et al, 2016). Replication timing (RT) also displayed high assortativity in the P-P network (RTAs=0.61), as expected by the excellent alignment between RT domains and chromatin interaction maps (Ryba et al, 2010; De and Michor, 2011; Boulos et al, 2015; for a visual representation of this phenomenon, see **Supp Figure 3B**).

**Figure 4.**
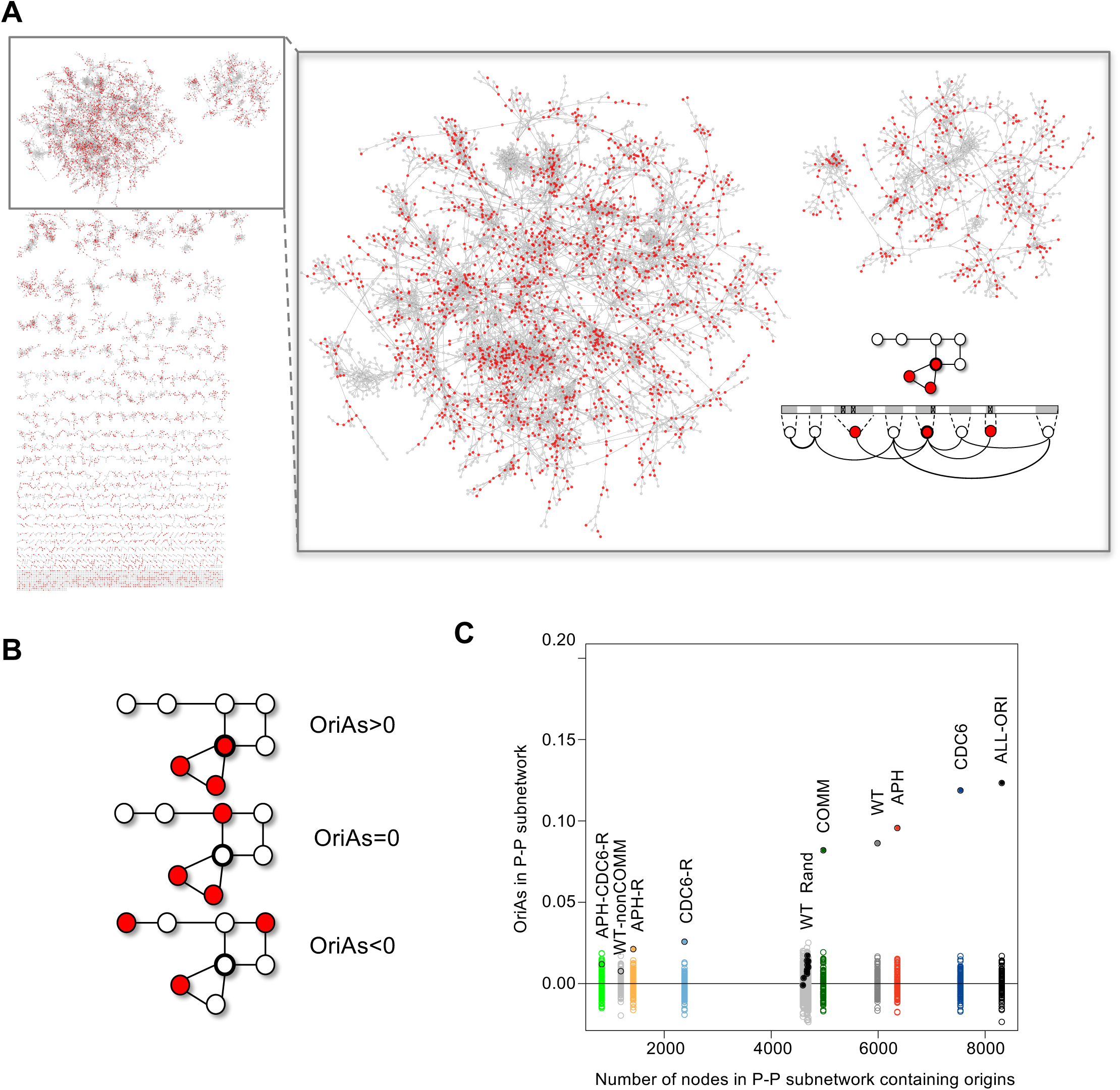
Mapping replication origins onto chromatin interaction networks. **A**. Panoramic view and detail of the mESC PCHiC subnetwork showing promoter-promoter (P-P) interactions with fragments containing WT origins marked in red. Each node (circle) represents a chromatin fragment bait. The link between two nodes is drawn if the two fragments interact in the PCHiC assay after ChICAGO filtering. The schematic (bottom right) depicts the correspondence between an 8-node subset of the network, centered on the node with thick border, and the interactions between the corresponding genomic regions (light gray areas). Red color indicates fragments that contain origins (marked as crosses in the linear DNA representation). **B**. Schematic representation of the possible values of origin assortativity (OriAs) in the network. **C**. Assortativity of origin presence (OriAs) of the indicated origin datasets was calculated on the P-P subnetwork and plotted against the total number of origin-containing promoter fragments in each dataset. Filled circles represent OriAs values for each dataset and empty circles show the values of OriAs obtained after 100 origin label permutations (**Supp Figure 4A**). The solid black circles (WT Rand) represent 20 sets of TSS distance-preserving randomizations applied to the WT dataset.

Assortativity of origin presence (OriAs) was positive in the P-P subnetwork for the WT, APH and CDC6 datasets, as well as for a collection of all combined origins (ALL-ORI) and the subset of COMM origins (**Figure 4C**), and to a much lesser extent for the responsive subsets. The significance of OriAs values was measured in relation to assortativity values produced by random origin label permutations within the network (empty circles around the horizontal axis in **Figure 4C;** note that this is different from randomizing origin positions in the genome. See below, and **Supp Figure 4A** for a schematic description of the permutation method). Importantly, positive OriAs is restricted to experimental origins, as it was not observed in randomized sets of genomic positions mimicking origins even when the distance to the TSS was preserved (black dots in **Figure 4C**). In addition, OriAs was strictly dependent on P-P interactions, as it was not significantly different from random permutations in the entire PCHiC network or the P-O subnetwork **(Supp Figure 4B-C).** These analyses indicate that, specifically within the P-P subnetwork, chromatin regions containing origins tend to interact with other regions that also contain origins.

### Origin connectivity correlates with efficiency and RT

An origin subnetwork termed ori-net (7,611 nodes and 7,791 edges) was defined by the PCHiC fragments that contained at least one origin in the WT, APH or CDC6 datasets, since virtually all of them displayed some activity in WT cells. All across ori-net, clusters of connecting origins formed hubs in 3D that resemble the type of organization expected at replication factories (**Figure 5A).** The majority of origin-origin contacts spanned 100 Kb to 1 Mb (mean 550 Kb), with 10% of them spanning >1 Mb and 2% of them spanning >10 Mb (**Figure 5B)**. Origin-origin interactions were mainly intra-chromosomal, albeit some contacts were detected between different chromosomes (**Supp Figure 4D**). Approximately 80% of ori-net interactions were established within the same topologically associated domain (TAD), while the rest reflected inter-TAD links. The median distance between contacting origin fragments within a TAD was 190 Kb.

**Figure 5.**
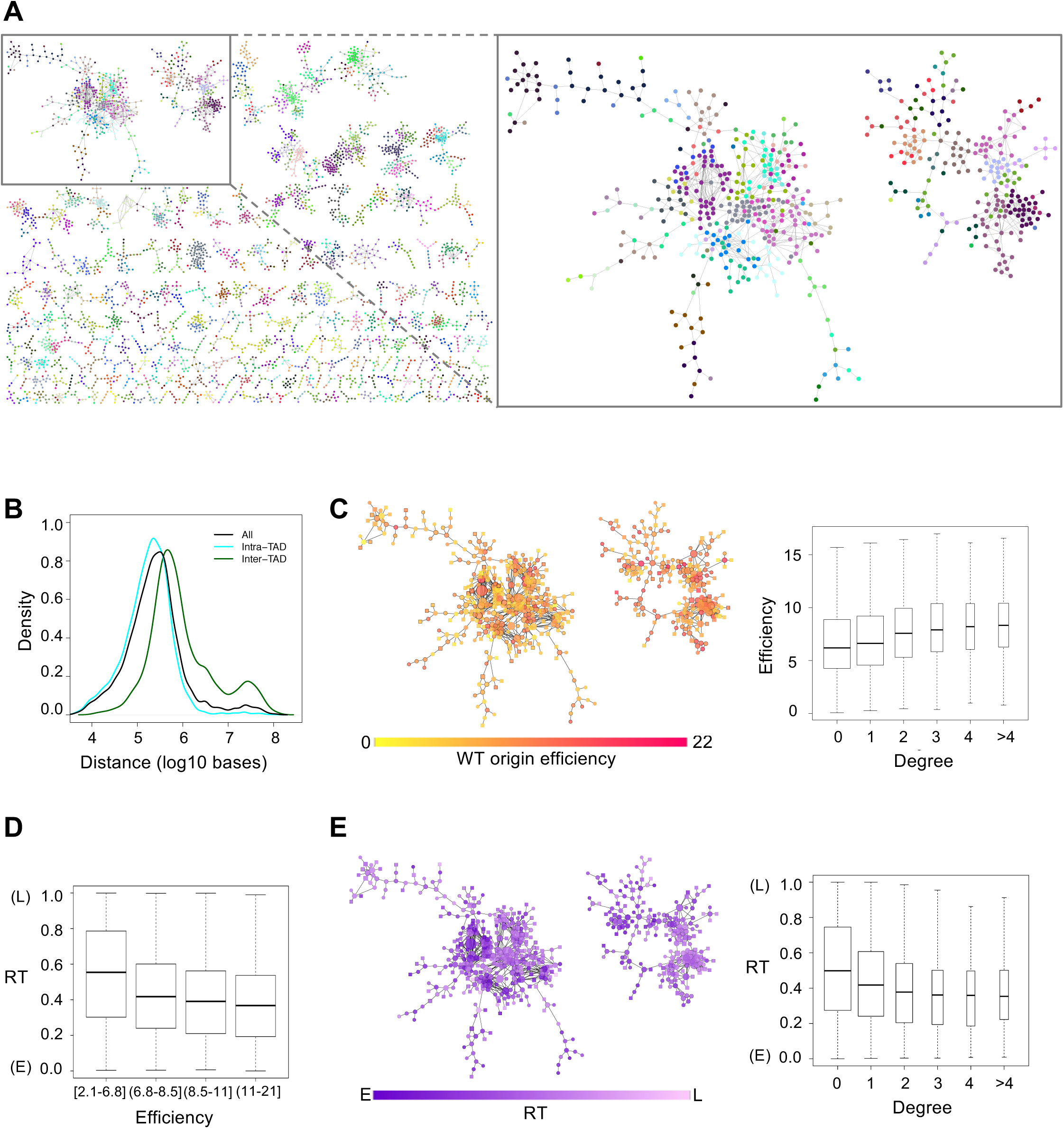
A 3D network of replication origins. **A**. Left, representation of a subset of ori-net (small, low-connected components not shown). Right panel shows the two largest connected components at higher magnification. Nodes (circles) represent genomic fragments containing origins. Connections between nodes are drawn when the fragments are connected in the PCHiC network. Node colour indicates the TAD in which the fragment is included. **B**. Distribution of distances spanned by contacts in the entire ori-net (black), the subset of connections linking origins in different TADs (dark green) and the subset of connections linking origins in the same TAD (cyan). **C**. Representation of the same ori-net components as in A (right), in which a yellow to red gradient in node colour represents increasing origin efficiency in each fragment. Box plots show the distribution of origin efficiency according to their connectivity in ori-net (degree; Pearson’s r=0.17, p<10^−16^). The efficiency of all origins was measured in the WT condition. **D**. Boxplots showing the correlation between RT and origin efficiency values (Pearson’s r=-0.25, p<10^−16^). **E**. Representation of the same ori-net components as above, in which a purple to pink gradient in node colour represents RT in each chromatin fragment. Box plots show the correlation (Pearson’s r=-0.18, p<10^−16^) between RT values and origin connectivity in ori-net. E, early; L, late replication.

A positive correlation between origin efficiency and node degree was observed, indicating that origins located in more connected nodes are activated more frequently in the population (**Figure 5C**). Besides ori-net, this effect was also detected when the individual WT, APH and CDC6 datasets were superimposed with the PCHiC network (**Supp Figure 5A**). In human and Drosophila cells, higher origin density and/or efficiency has been reported in early-replicating chromosomal domains (Cadoret et al, 2008; Besnard et al, 2012; Lubelsky et al, 2014; Cayrou et al, 2015). A cross-analysis of our origin datasets stratified by efficiency with published RT data for mESCs (Hiratani et al, 2010) also revealed a direct correlation between efficiency and RT, in which more efficient origins tend to be early-replicating (**Figure 5D**). Therefore, a positive correlation between origin-origin connectivity and RT can be predicted. Indeed, origins located at nodes with higher degree displayed earlier RT than those located at lower-degree nodes (**Figure 5E** and **Supp Figure 5B**). Taken together, these results indicate that origins at highly connected nodes in a promoter-centered 3D origin interaction network tend to activate early in S phase and display higher efficiency in the population.

### Short- and long-range interacting origins display similar efficiency

To further test the link between origin connectivity and frequency of activation, the assortativity of origin efficiency (OriEfAs) was calculated. Positive OriEfAs was observed for WT, CDC6, APH and COMM origins in the P-P subnetwork, indicating that origins that interact with each other have a tendency to be activated with similar efficiency. The highest OriEfAs value was obtained with a combined dataset of origins from the WT, APH and CDC6 sets (ALL-ORI), in which their efficiency was taken from WT conditions. In contrast, OriEfAs values in responsive origins were close to those obtained in random permutations of origin efficiency (**Figure 6A** and **Supp Figure 5C**).

**Figure 6:**
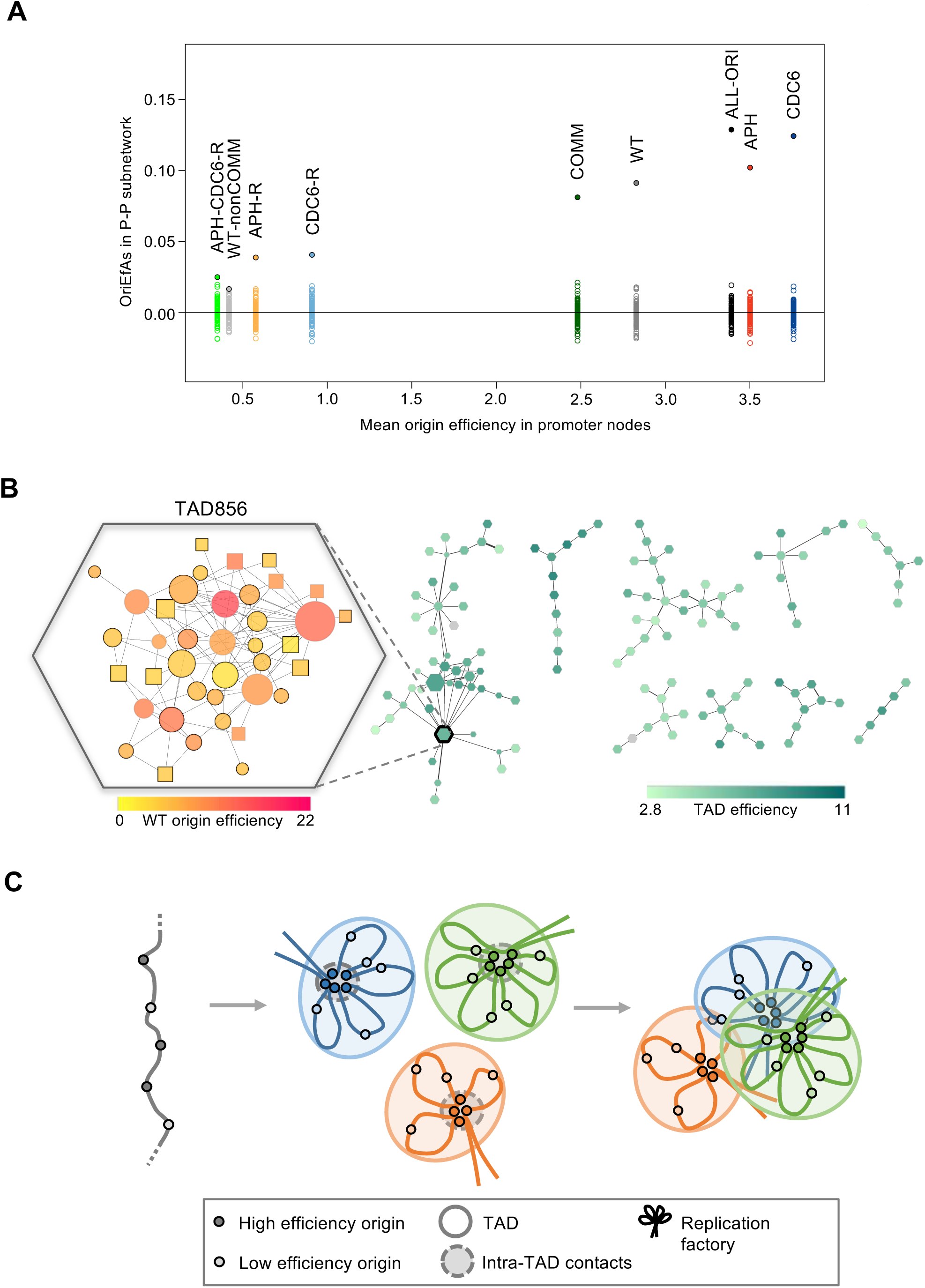
Interacting origins tend to be activated with similar efficiency. **A**. Assortativity of origin efficiency (OriEfAs) in the indicated datasets measured in the P-P subnetwork and plotted against the average efficiency of origins in the P nodes in each dataset. Filled circles represent OriEfAs values for each dataset and empty circles show the values of OriEfAs obtained after 100 efficiency permutations (**Supp Figure 5C**). **B.** Subset of a TAD interaction network with nodes (hexagons), in which a light to dark green gradient in node color represents increasing average efficiency of the origins in each TAD. The left inset shows the origin interaction network within TAD856. Circles indicate P nodes and squares indicate O nodes. Fragments containing COMM origins are marked with a thick border. Node color represents origin efficiency (measured in WT conditions), and node size is proportional to the number of connections within the TAD. **C.** Schematic model of the hierarchical organization of replication origins. Single replication origins are close to each other at the base of chromatin loops forming replication factories, in turn brought into 3D contact across longer genomic distances even spanning multiple TADs. Origins brought in 3D contact have similar efficiency values and RT.

A visual representation of these phenomena is presented in the chromosomal domain corresponding to TAD856, highlighted in **Figure 6B**. Origins located at P and O fragments have been plotted as circles and squares, respectively. The size of each node is proportional to its number of connections, and color intensity indicates average efficiency. With this representation, it becomes apparent that origins located at promoters (circles) tend to be more connected than origins located at other ends (squares), and larger, more connected nodes tend to be darker, indicating that they contain more efficient origins. Interestingly, efficiency was also assortative in a TAD-level contact network in which nodes correspond to entire TADs (TADOriEf = 0.27, random values in the −0.04-0.04 range). This effect is visualized in **Figure 6B** (right), in which TADs are colored according to the average efficiency of the origins contained in them. We also observed that origins belonging to different TADs that interact in 3D display similar RT **(Supp Figure 5D)**. These data suggest that interacting origins tend to replicate synchronously, regardless of whether they belong to the same TAD or separate TADs, and are activated with similar efficiencies across the population.

### Origin integration in alternative chromatin contact networks

Finally, origin assortativity was measured in alternative chromatin contact networks generated from RNA polymerase II and SMC1 ChIA-PET datasets (Zhang et al, 2013; Dowen et al, 2014) as well as a haploid mESC HiC dataset (Stevens et al, 2017). Despite the fact that these networks displayed very different levels of origin coverage, long-range interactions and connectivity, WT, APH, CDC6 and COMM datasets showed significantly higher assortativity than expected at random in all of them (**Supp Figure 6**). In these cases, assortativity was also observed in some of the responsive origin subsets, probably reflecting the fact that these networks include many contacts between non-promoter regions.

## DISCUSSION

The activation of ‘dormant’ replication origins in response to stress is a crucial mechanism to maintain genomic stability (Ge et al, 2007; Ibarra et al, 2008, Doksani et al, 2009). However, the genomic positions and characteristics of stress-responsive origins are only now starting to be elucidated. In this study, we have compared the activity of replication origins in control mESCs or under situations of mild replicative stress induced by aphidicolin or CDC6 ectopic expression. The distribution and enrichment of mESC origins at different genomic elements were remarkably similar in WT, APH and CDC6 conditions. Since the original report that initiation of mammalian DNA replication occurred frequently at CGIs (Delgado et al, 1998), most origin mapping efforts have revealed a strong association of origins with gene promoters. Our extensive cross-analyses of WT, APH and CDC6 origins with a large collection of chromatin marks underscore this connection. Accordingly, a recent genome-wide mapping of origins in chicken DT40 cells has uncovered that the fraction of mammalian origins functionally conserved in avian cells is 2-3-fold higher among those associated with CGI and TSS than in the rest (Massip et al, 2019).

The intersection of origin datasets, as defined by a strict combination of two distinct peak-calling algorithms, hinted at the existence of many stress-responsive origins in the mouse genome. Interestingly, the distribution of WT SNS-Seq reads around these origins revealed that they were also active in a fraction of the cell population undergoing an unchallenged S-phase. This pattern is better reconciled with a stochastic rather than deterministic model of origin usage. The term ‘stochastic’ should not be understood as random initiation from any genomic position; it rather indicates that, while origins are located at preferred sites, not every cell activates exactly the same set (**Figure 2A**). In this view, stress-responsive origins correspond to origins that have remained silent in a given cell but might have been activated in other cells in the population. Deterministic models may seem more intuitive, but stochastic origin activation has been described in unicellular yeasts (Patel et al, 2006; Czajkowsky et al, 2008; Rhind et al, 2013) and is compatible with the completion of DNA replication following a replication timing program in mammalian cells (reviewed by Rhind and Gilbert, 2013).

We observed that DNA polymerase slowdown, or increased levels of the licensing protein CDC6, caused a global increase in origin activity. This effect was observed at the population level, but not necessarily at every origin (see schematic in **Figure 2G**). Of note, a recent report has shown that the most efficient origins in human cells can be further stimulated upon mild hydroxyurea treatment (Chen et al, 2019). This study and ours have reached compatible conclusions about stress-responsive origin activation, despite having used different cell types, origin mapping methods and efficiency calculation algorithms.

The stratification of mESC origins according to their firing efficiency, i.e. their frequency of activation in the population, indicated that the most efficient origins were those colocalizing with CGI, TSS and chromatin features characteristic of promoter elements. This fact likely explains that these origins have been frequently identified, regardless of the mapping method used. For instance, recent studies using either Oka-Seq or MCM7 chromatin immunoprecipitation in human cells have located the preferred initiation sites immediately upstream of the TSS (Sugimoto et al, 2018; Chen et al, 2019). A detailed analysis of firing efficiency, gene length and RNA polymerase II (RNAPII) occupancy revealed a strong positive correlation between origin activity and TSS of genes with high RNAPII occupancy, likely ensuring co-orientation of replication with the most highly transcribed regions of the genome (Chen et al, 2019). In turn, the marks that characterize the elongating form of RNAPII, such as phosphorylation of its CTD Ser2 and the presence of H3K36me2/3 modifications, are amongst the least enriched features at origins. The lower abundance of active origins at regions where transcription elongation takes place may have evolved to avoid or minimize transcription-replication conflicts.

Our study revealed that the frequency of origin activation is also influenced by the three-dimensional organization of the genome, particularly by the number of contacts established between chromatin fragments containing origins. The impact of higher-order chromatin organization on DNA replication has been established at the level of chromatin TADs, which correlate almost exactly with replication timing domains (Pope et al, 2014). However, how it affects the relative positions and activation of individual origins was poorly understood. The integration of linear origin maps in chromatin interaction networks strongly supports the proposed architectural organization of origins in DNA replication factories, a still speculative but widely-accepted model for replication units in which local clusters of origins are arranged in physical proximity while the stretches of inter-origin DNA are looped out (**Figure 6C**; Hozak et al, 1993; Jackson and Pombo, 1998; Berezney et al, 2000). This architectural arrangement is facilitated by the cohesin complex (Guillou et al, 2010) and likely creates a favorable environment for the local concentration of origin-binding and origin-activating proteins. Most replication factories are likely located within single TADs, but a fraction of the origin-origin contacts spanned much longer distances that reflect inter-TAD interactions (**Figure 6C**), likely belonging to the same chromatin compartment. In the latter case, the presence of origins at both ends correlated strongly with both fragments having a similar RT and origin activation efficiency.

Based on previous work and the new analyses reported in this study, we postulate that within each factory, highly efficient origins are located at the bases of the DNA loops, establishing multiple connections between them and probably benefiting from the local accumulation of origin-binding proteins and activating factors. In contrast, dormant origins would be preferentially located outside the factory core, establishing fewer or no connections with other origins and therefore being activated with lower frequency (**Figure 6C)**. In normal conditions, most of these origins will be passively replicated from the forks derived from the core. Upon stalling of these forks, however, less connected origins increase their frequency of activation. It remains to be determined whether they fire at their original location or need to be relocated three-dimensionally to the factory core.

It can be speculated that replication factories display characteristics similar to phase-separated multimolecular assemblies such as Cajal bodies, nuclear speckles or superenhancers (Hnisz et al, 2017). These membrane-less structures are formed in discrete nuclear zones with a high density of proteins and nucleic acids that establish cooperative interactions between them. For instance, a recent study has revealed how a group of subtelomeric origins in fission yeast are tethered by shelterin components to a local domain enriched in Rif1 and protein phosphatase 1, imposing late replication (Ogawa et al, 2018). The proteins that recognize replication origins, ORC and CDC6, also serve as molecular chaperones capable of attracting and assembling many additional proteins to the factory core, including CDT1, MCM, CDC45, GINS, PCNA, and DNA polymerases that establish multiple contacts between them. We have noticed that human ORC1 is predicted to contain a large intrinsically disordered region (IDR; as analyzed by PONDR software, www.pondr.com). The presence of IDRs is a common feature in proteins that facilitate multivalent interactions and higher-order signaling assemblies (reviewed by Wright and Dyson, 2015). The bursts of DNA synthesis created by the activation of several adjacent origins, classically visualized as discrete nuclear BrdU or PCNA foci, would parallel the bursts of transcription driven by superenhancers.

While many structural and mechanistic aspects of mammalian replication factories remain to be elucidated, it is likely that replicating chromatin also forms specific domains whose segregation into specific areas of the nucleus may play a role in the known connections between RT, accumulation of mutations and copy number variations that determine the evolvability of specific genomic regions (Juan et al, 2014; Wu et al, 2018).

## MATERIALS AND METHODS

### Cell lines and culture

TetO-CDC6 mouse embryonic stem cells (mESCs) were derived from the TetO-CDC6 mouse model (Muñoz et al., 2017). TetO-CDC6 mESCs were cultured on 0.1% gelatin-coated plates in Dulbecco’s modified Eagle’s medium (DMEM) with Ultraglutamine 1 and 4.5 g/L glucose (Lonza) supplemented with 15% FBS (Sigma), 50 U/mL Penicillin - 50 mg/mL Streptomycin (Invitrogen), Minimum Essential Medium Non-Essential Aminoacids (MEM NEA; Invitrogen), 100 μM 2-mercaptoethanol (Invitrogen) and 10^3^ U/mL ESGRO mLIF Medium Supplement (Millipore). To induce CDC6 overexpression, 1 µg/ml doxycycline (dox, Sigma) was added to the medium for 30 h. To induce mild replication stress, cells were treated with 0.5 µM aphidicolin (Sigma-Aldrich) for 2.5 h.

### Flow cytometry

To monitor DNA content, cells were stained overnight with 50 μg/ml propidium iodide (PI; Sigma-Aldrich) in the presence of RNase A (10 μg/ml, Qiagen). In order to analyse DNA synthesis, cells were pulse-labelled with 20 μM BrdU for 30 min, trypsinized, washed in PBS and fixed with −20°C 70% ethanol for 24 h. 2 M HCl was added for 20 min at RT, before washing cells twice with PBS and incubating in blocking solution (1% bovine serum albumin in PBS, 0.05% Tween-20) for 15 min at RT. FITC-conjugated anti-BrdU antibody (BD Biosciences Pharmigen) was added for 1 h at 37°C. Samples were analyzed in a FACS Canto II cytometer (Becton-Dickinson) and data was processed with FlowJo V 9.4 or V.10.1 (Three Star).

### Analysis of DNA replication in stretched DNA fibers

Cells were pulse-labelled sequentially with 50 μM CldU (20 min) and 250 μM IdU (20 min), harvested and resuspended in PBS (0.5 x 10^6^ cell/ml). 2 μl drops of cell suspension were placed on microscope slides and lysed with 0.5% SDS, 0.2 M Tris pH 7.4, 50 μM EDTA in 10 μl for 6 min at RT. Slides were tilted 15 degrees to spread DNA fibers, air-dried, fixed in −20 C methanol:acetic acid (3:1) for 2 min and stored at 4 C overnight. Slides were then incubated in 2.5 M HCl (30 min/ RT) to denature DNA and washed (3x) in PBS. Blocking solution (1% bovine serum albumin in PBS, 0.1% Triton X-100) was added for 1 h at RT. Slides were incubated with anti-CldU, IdU and ssDNA primary antibodies for 1h at RT, washed and incubated with the corresponding secondary antibodies for 30 min. Prolong mounting media (Invitrogen) was used. Images were acquired in a DM6000 B Leica microscope with an HCX PL APO 40x, 0.75 NA objective. Fork rate values were derived from the length of IdU tracks, measured using ImageJ software, and a conversion factor of 1 μm = 2.59 kb (Jackson and Pombo, 1998). >300 tracks were measured per condition. Three biological replicates of each experiment were performed.

### Short Nascent Strand purification

10^8^ growing cells were harvested and lysed for 15 min in 0.5% SDS, 50 mM Tris pH 8.0, 10 mM EDTA and 10 mM NaCl. 100 μg/ml Proteinase K (Roche) was added and samples were incubated overnight at 37°C. DNA was isolated by standard phenol purification and EtOH precipitation, resuspended in TE (10 mM Tris pH 8.0, 1 mM EDTA) supplemented with 0.1 U/μl RNAseOUT (Invitrogen), and stored at 4°C for at least 48 h. Following heat denaturation (100°C/ 10 min), DNA samples were loaded onto 5-20% sucrose gradients and fractionated according to size by centrifugation (SW-40Ti rotor; Beckman Coulter Optima L-100 XP; 20 h/ 78000 rcf / 20°C) as described (Gómez and Antequera, 2008). DNA from approximately 13 x 1-ml fractions was precipitated with ethanol and analysed in 1% alkaline agarose gel electrophoresis. Fractions 4-5, corresponding to DNA fragments of 300-1500 nucleotides, were selected for further analysis. For each experiment, DNA samples were pooled from two gradients.

DNA samples were treated with 100 U of T4 polynucleotide kinase (PNK, Thermo Fisher) in the presence of 1 mM dATP (Roche) and 40 U of RNAseOUT (Thermo Scientific) for 30 min at 37°C. The PNK reaction was stopped by the addition of 6.25 µg proteinase K, 0.125% sarkosyl and 2.5 μM EDTA (30 min/ 37°C). Samples were heat-denatured (95°C/ 5 min) and incubated o/n at 37°C with 150 U of λ-exonuclease (λ-exo; Thermo Scientific) in the presence of 40 U of RNAseOUT. Reactions were heat-inactivated (75°C/ 10 min) and DNA was recovered by EtOH precipitation. To increase the purity of SNS, three cycles of PNK treatment and λ-exo digestion were performed. Each digestion step was controlled by adding 50 ng of linearized pFRT-myc plasmid (kindly provided by Dr. Susan Gerbi, Brown University, USA) to 5% of the digestion reaction and incubated in the same conditions. pFRT-myc contains two G-quadruplex-forming sequences, reported to be digested less efficiently by λ-exo (Foulk et al, 2015). Control λ-exo reactions were analysed in 1% agarose gels to confirm full digestion of the plasmid. To control for SNS enrichment at the Mecp2 origin, qPCR reactions were performed in duplicates using ABI Prism 7900HT Detection System (Applied Biosystems) and HotStarTaq DNA polymerase (Qiagen) according to manufacturer’s instructions. Data was analysed in Applied Biosystems Software SDS v2.4. Primer sequences for Mecp2 origin and Mecp2 flanking region are indicated in **Supp Table 3**.

### SNS-Seq library preparation and high-throughput sequencing

RNA primers were removed with RNAse A/T1 Mix (Roche) for 60 min at 37 °C. 100 μg/ml Proteinase K was added (30 min, 37°C) and DNA was extracted and precipitated. ssDNA was converted to dsDNA using 50 pmol of random hexamer primers phosphate (Roche) as previously described (Cadoret et al, 2008). Primer extension was performed by incubation with 10 mM dNTPs (Roche) and 5 U exo-Klenow Fragment (New England Biolabs) for 1 h at 37 C followed by incubation with 80 U of TaqDNA ligase (New England Biolabs; 50 C/ 30 min). DNA was extracted, precipitated and resuspended in TE. For the input sample, 4 x 10^7^ mESCs were lysed in 1 % SDS, 50 mM Tris-HCl pH 8.0, 10 mM EDTA (2 x 10^7^ cell/ml) and sonicated in a Bioruptor device (Diagenode) for 25 min at 30 s intervals. DNA was extracted with phenol/chloroform, ethanol-precipitated and resuspended in 0.5x TE. DNA libraries were prepared at the Fundación Parque Científico de Madrid (FPCM) using NEBNext® Ultra™ II DNA Library Prep Kit for Illumina (New England Biolabs) and purified with Agencourt AMPure XP beads (Beckman Coulter). Each library was sequenced using single-end 75 bp reads (120-140 x 10^6^ reads per sample) in a NextSeq500 System (Illumina).

### Whole cell extract preparation and immunoblots

Cells were harvested and resuspended in Laemmli sample buffer (50 mM Tris-HCl pH 6.8, 10% glycerol, 3% SDS, 0.006% w/v bromophenol blue, 5% 2-mercaptoethanol) at 10^6^ cells/ml. Extracts were sonicated for 30 s at 15% amplitude in a Branson Digital Sonifier. Standard protocols were used for SDS-polyacrylamide gel electrophoresis and immunoblotting. Primary antibodies used in this study are listed in **Supp Table 4**. Horseradish peroxidase (HRP)-conjugated secondary antibodies (GE Healthcare) and ECL developing reagent (Amersham Biosciencies) were used.

## COMPUTATIONAL METHODS

### SNS-seq data analysis

The quality of sequencing reads was analysed with FastQC (www.bioinformatics.babraham.ac.uk/projects/fastqc/). Short sequences, adaptor sequences and read duplicates were removed. Reads were analysed with the RUbioSeq pipeline v3.8 (Rubio-Camarillo et al., 2017) using BWA v0.7.10 (Li and Durbin, 2009), SAMtools v0.1.19 (Li et al., 2009), Picard tools v1.107 (broadinstitute.github.io/picard/) and MACS v2.0.10 (Feng et al., 2012), and the GRCm38/mm10 mouse reference genome. When MACS was used, peak calling was performed versus input. When the algorithm described in Picard et al (2014) was used, genome segmentation was based on RT data that accurately matches the read coverage differences between segments. Common peaks were obtained using BedTools v2.23.0 (Quinlan, 2014) with parameters: -f 0.1 -r -wa ‒u. For common peaks between MACS and Picard the genomic coordinates defined by Picard were used in additional analyses. Peaks located on chromosome Y were excluded from the analysis. The SNS-seq WT I dataset, taken from Almeida et al (2018; GSE99741), was generated in parallel to all other SNS-seq samples, which were specifically designed for this study. Read distribution around peak centres was generated using seqMINER v1.3.3e.

### Epigenomic features and chromatin states analyses

Genomic coordinates of origins were converted from mm10 to mm9 genome assembly with LiftOver (https://genome.ucsc.edu/util.html). Origins were intersected with the genomic features depicted in **Supp Table 5** or with a set of previously compiled epigenomic features (Juan et al, 2016). Each epigenomic feature dataset was ‘discretized’ in 200 bp windows: the presence of a given mark within a 200 bp window was scored as 1, and its absence as 0. The overlap between origin fragments and the genomic windows was calculated using findOverlaps in the genomicRanges R package. The number of origins overlapping with each feature was calculated for the experimental sets of origins and for 1,000 sets of origins randomly shuffled along the genome (excluding low-mappability regions). The enrichment of origins at any particular feature was calculated as the ratio between the number of origins overlapping the feature and the median of all randomizations, generating an empirical p-value. In the calculations of origin enrichment at chromatin states, some of the 20 states defined by Juan et al (2016) corresponding to similar chromatin functions were merged according to the definitions indicated in **Supp Table 6**.

### Analysis of origin efficiency

In each sample, the efficiency of each individual origin was determined following three steps: (1) The sum of reads covering each nucleotide (i.e. the sum of per base read depths) of the origin was calculated using samtools function bedcov with default parameters. This analysis was done after downsampling every dataset to the lowest coverage obtained (APH-I). Down-samplings were performed a total of 10 times, and the median value was determined for each origin; (2) Background was calculated for each sample as the sum of per base reads in random genomic fragments that are not origins but have the same size. After 100 randomized relocations, a function of median background noise by size of the fragment was calculated. The estimated background noise for each origin was subtracted from the initial values; (3) Background-corrected values were normalized dividing by the size of the origin. Finally, mean efficiencies of origins from biological replicates were calculated as an average of efficiency between the two replicates.

### Origin set randomization

Randomized sets of origins were obtained by relocating all origins in a different position choosing from the whole genome, excluding low-mappability regions and ensuring that randomized origins did not overlap with real ones. For the TSS distance-preserving randomization used in the network analysis, random origins were placed at new genomic positions but maintaining the same distance from a TSS as the real ones. In this process, some candidate random origins were placed at new locations with the correct distance from the target TSS, but accidentally closer to another TSS. These candidate random origins were discarded and randomized again. If after 1,000 randomization attempts, some origins (always < 2%) still failed to match the randomization criteria, the distance from the TSS was progressively increased until these origins were successfully relocated. Custom scripts can be found at https://github.com/VeraPancaldiLab/RepOri3D.

### Low-mappability regions

The scanquantile peak-calling algorithm (Picard et al, 2014) was used on a sequenced genomic DNA input, with the same parameters set for the SNS-seq samples. The resulting peaks, as well as the small gaps present in the genome segmentation needed for peak-calling, were marked as non-mappable regions (ShadeAreas files at https://github.com/VeraPancaldi/RepOri3D). Subtelomeric and pericentromeric regions, extended from the UCSC database telomere annotation by visual inspection in the browser, were also added.

### Integration with chromatin interaction maps

Linear origin maps were integrated into the following 3D chromatin interaction maps for mESCs: Promoter-Capture HiC (PCHiC; Schoenfelder et al, 2015), SMC1 ChIA-PET (Dowen et al, 2014), RNA Polymerase II (RNAPII) ChIA-PET (Zhang et al, 2013) and haploid mESC HiC (Stevens et al, 2017). PCHiC and ChIA-PET networks were processed as described in Pancaldi et al (2016). The PCHiC contact map was processed using the CHiCAGO pipeline, which identifies significant 3D contacts starting from the raw Capture HiC data. The other networks were generated starting from the contacts provided in the original references. Origin positions were mapped to the chromatin fragments of 3D maps using findOverlaps (genomicRanges package). Origin efficiency of the chromatin fragment was calculated as the mean of the efficiencies of origins that overlap with this fragment.

### Network analysis

All network analyses including correlations between degree, efficiency and replication timing (RT) were performed using the igraph package in R and standard R functions. TAD definitions were taken from Dixon et al (2012). Networks were visualised using Cytoscape v3.6.1. Replication timing data for three mESC cell lines was downloaded from Hiratani et al (2010) and combined in our analysis. The median value for probes overlapping each origin was taken and individual origins were ranked based on RT in the range 0 to 1 (early to late). When indicated, the probes within each chromatin fragment in the 3D chromatin network were combined to give an average RT value for the entire fragments.

### Assortativity analysis

Origin Assortativity (OriAs) and Assortativity of Origin Efficiency (OriEfAs) were calculated using the previously described measure of Chromatin Assortativity (Pancaldi et al, 2016). Briefly, assortativity is defined as the Pearson correlation coefficient of the presence of an origin (OriAs) or of the value of origin efficiency (OriEfAs) across all pairs of nodes that are connected with each other. This value was calculated with the assortativity function in the igraph package for R. Chromatin assortativity (ChAs) for a particular feature is analyzed in relation to the abundance of said feature. For example, if a particular mark is found in the majority of the fragments in the network, its localization in specific areas of the network cannot be observed and the value of ChAs will be low. On the contrary, if a certain feature is detected only in a small subset of fragments, but they interact preferentially with each other, the ChAs measure will be high. Despite the fact that assortativity is better defined on continuous values than on binary ones, we found OriAs to be very similar to OriEfAs.

### Statistical analysis

Statistical analyses relative to the wet-lab experiments were performed using Prism v4.0 (GraphPad Software) or Microsoft Excel v15.38. For comparison of two data groups, two-tailed paired Student’s t-test was used. In the analysis of fork rate in stretched DNA fibers, a nonparametric Mann-Whitney rank sum test was used. To statistically assess the overlap between replication origins and genomic features or chromatin marks, we compared the number of overlaps between origins and features to the number of overlaps calculated after randomly relocating the origins in the genome 1,000 times. Empirical p-values were obtained as (*r*+1)/(*n*+1), where *n* is the number of randomizations and *r* is the number of randomizations that produce a test statistic greater (or smaller) than or equal to that calculated for the real data. All statistical analyses for the computational part were performed using R. Scripts and R notebooks used to analyze origins in their 3D context can be found on https://github.com/VeraPancaldiLab/RepOri3D.

## ACKNOWLEDGEMENTS

JM and MG thank all members of our laboratories for useful discussions. We are grateful to Ana Cuadrado, Daniel Giménez and Ana Losada (CNIO, Madrid), Francisco Antequera (IBFG, Salamanca), Biola Javierre (Josep Carreras Leukaemia Research Institute, Barcelona) and Raphael Mourad (Paul Sabatier University, Toulouse) for comments on the manuscript. We thank Francisco Sanchez-Rivera (MSKCC, New York) for discussions about phase-separated assemblies, and Luis Quintales (IBFG, Salamanca, Spain) for computational help and advice in the initial stages of the project. The work was supported by grants BFU2013-49153-P and BFU2016-80402-R to JM and BFU2016-78849-P to MG (Spanish Ministry of Science, Innovation and Universities, co-funded with European Union ERDF funds). Additional support was provided by a “CNIO Friends” postdoctoral fellowship and the Chair of Bioinformatics in oncology of the CRCT (VP) as well as predoctoral Fellowships from CNIO-La Caixa (KJ and MR), the Portuguese Foundation for Science and Technology (FCT-SFRH/BD/81027/2011 to RA), and the Spanish Ministry of Science, Innovation and Universities (BES-2014-070050 to JMF-J).

## Notes

#### Summary of Updates

Incorrect reference to Figure 6D amended to reference figure 6C.

https://github.com/VeraPancaldiLab/RepOri3D

