## Supplementary Tables and Figures for "Three-dimensional connectivity and chromatin environment mediate the activation efficiency of mammalian DNA replication origins"

by Jodkowska, Pancaldi *et al*

**CONTENTS**


1. Supplementary Figure Legends 1-6
2. Supplementary Tables 1-6
3. Supplementary references


#### 1. Supplementary Figure Legends

**Supplementary Figure 1 (related to Main Figure 1). Origin identification by SNS-Seq. A**. Schematic of the SNS-Seq protocol and examples of DNA size fractionation by sucrose gradient centrifugation. Red boxes indicate the fractions collected in each case. **B.** Left, flow cytometry plots of BrdU incorporation in mESCs in the absence or presence of aphidicolin (top) or Cdc6 overexpression (bottom). The percentage of cells in G1, S or G2/M phases are indicated. Center, quantification of fork progression rate in the same cells (***$p<0.001$ in Mann-Whitney test). Right, immunoblots showing the indicated markers of DNA damage (top) and the level of overexpression of CDC6 protein (bottom). Mek2 and Ponceau-S are shown as loading controls. **C.** A plasmid containing two G4-forming sequences (Pu27, Pu30) was linearized with BglII and mixed with SNS preparations before λ-exo digestion to assess the completeness of the enzymatic reaction. After three rounds of digestion, no exonuclease-resistant bands derived from the plasmid were detected (compare to no λ-exo reaction; last lane). **D.** Representative SNS profile and qPCR enrichment of WT, APH and CDC6 SNS samples around the origin Mecp2 and a flanking (F) region located 885 bp downstream. The position of a CGI at the Mecp2 locus is indicated (green bar).

**Supplementary Figure 2 (related to Main Figures 2 & 3). Enrichment of origin datasets at genomic regions with different epigenetic marks. A.** Histograms show the enrichment of WT, APH and CDC6 origins in genomic regions carrying the indicated epigenetic marks. All enrichments are significant at $p<0.005$ (empirical p-value, 1000 randomizations, see Methods). **B.** Genome browser screenshots of 16 examples of APH-R, CDC6-R and APH+CDC6-R origins in which an enrichment in SNS reads is also detected in WT cells. See text for details. **C.** Histograms show the enrichment of WT, APH and CDC6 origins, classified in 4 groups according to their relative efficiency, in genomic regions carrying the indicated epigenetic marks. All enrichments are significant at $p<0.05$ except for the ones marked as n.s. (not significant).

**Supplementary Figure 3 (related to Main Figure 4). A. Visualization of origins in the PCHiC network.** Red nodes represent fragments containing WT origins in the largest connected component of PCHiC. **B**. **Visualization of RT values projected on the largest connected components of the P-P subnetwork.** Node color represents RT from dark (early) to light purple (late).

**Supplementary Figure 4 (related to Main Figures 4 & 5). Origins are neither assortative in the PCHiC network nor in the P-O subnetwork. A.** Origin label permutation procedure. Left, schematic representation of the linear map and corresponding network of a subset of PCHiC. Interacting chromatin fragments are represented by grey regions. Crosses indicate detected origins. In the network, chromatin fragments are represented as circles, which are red if containing at least one origin. Right, permutations are performed by shuffling the origin label across the network nodes, maintaining the number of red nodes. **B**. OriAs of the indicated origin datasets in the entire PCHiC network. **C.** Same as (B), in the PCHiC P-O subnetwork. **D. The majority of origin-origin interactions are intrachromosomal.** Representation of the PCHiC origin subnetwork (ori-net), in which nodes are colored by chromosome.

**Supplementary Figure 5 (related to Main Figures 5 & 6). More connected origins are more efficient and replicate earlier. A.** Correlation between origin efficiency and node degree in WT, APH and CDC6 datasets. **B.** Correlation between RT and node degree in WT, APH, CDC6 datasets. **C.** Schematic of the origin efficiency permutation procedure. Top, a linear map containing 8 interacting fragments (labeled A-H) and its corresponding network representation is shown. Fragments are represented by grey regions, and crosses indicate detected origins. In the network, chromatin fragments are represented as circles, in which the color represents the average efficiency of the origins contained in them: red, high; orange, intermediate; yellow, low. White fragments contain no origins. Bottom, origin efficiency permutation performed by shuffling the efficiency values of network fragments. **D.** Distribution of RT differences between interacting PCHiC fragments containing origins. For each pair of nodes, the difference of their RT values was calculated. Node pairs corresponding to intra- and inter-TAD contacts were evaluated separately. As a control, the distribution of RT differences between random pairs of fragments (not PCHiC edges) was also calculated. Black horizontal lines represent the median difference in RT for each category of pairs of nodes.

**Supplementary Figure 6 (related to Main Figures 4-6). Origin efficiency assortativity (OriEfAs) in alternative chromatin networks. A.** RNA polymerase II ChIA-PET chromatin network. **B**. SMC1 ChIA-PET chromatin network **C.** HiC chromatin network.

#### 2. Supplementary Tables

| Origin set | Total origins | Origins overlapping with PCHiC fragments |
|---|---|---|
| WT | 20,174 | 7,886 |
| APH | 31,685 | 8,996 |
| CDC6 | 31,402 | 10,608 |
| COMM | 11,998 | 6,028 |
| APH-R | 17,310 | 2,548 |
| CDC6-R | 17,676 | 3,878 |
| APH+CDC6-R | 8,269 | 1,283 |
| WT-nonCOMM | 8,176 | 1,858 |
| ALL-ORI | 46,988 | 13,038 |

**Supplementary Table 1.** Total number of origins *(mm9 coordinates)* in the indicated subsets and number of origins that are located at PCHiC nodes**.**

| | WT | Origin randomization (mean) | TSS distance-preserving origin randomization (mean) |
|---|---|---|---|
| PCHiC | 7886 | 2655 | 6812 |
| P nodes | 6455 | 1201 | 5527 |
| O nodes | 1587 | 1498 | 1501 |
| PP subnetwork | 5109 | 894 | 4308 |
| PO subnetwork | 6917 | 2451 | 5923 |
| Total origins | 20174 | 20328 | 20345 |
| proportion of origins overlapping PCHiC | 0.39 | 0.13 | 0.33 |
| proportion of origins overlapping P nodes | 0.32 | 0.06 | 0.27 |
| proportion of origins overlapping O nodes | 0.080 | 0.07 | 0.07 |
| proportion of origins overlapping P-P fragments | 0.25 | 0.04 | 0.21 |
| proportion of origins overlapping P-O fragments | 0.34 | 0.12 | 0.29 |

**Supplementary Table 2.** Overlap of WT origins and randomized origins (preserving or not the distance to the TSS) with PCHiC, P and O nodes and PP, PO subnetworks.

| Gene/region | primer name | primer sequence 5'- 3' |
|---|---|---|
| Mecp2 origin | Mecp2 43 Fwd | CTACCCGCCCCCCAGCAAG |
| | Mecp2 44 Rv | GTGAGTGGGACCGCCAAGG |
| Mecp2 flank | Mecp2 1 Fwd | GCATCCAATGCTCTTTGTGC |
| | Mecp2 14 Rv | GTCTCTTGTTGAGCATTTGT |

**Supplementary Table 3. Primers used for qPCR**.

| Antibody | Use | Supplier | Ref/Catalogue # | Species |
|---|---|---|---|---|
| CDC6 | WB | Millipore | 05-550 | Mouse |
| p53-P-S15 | WB | Cell Signalling | 9284S | Rabbit |
| p53 | WB | Cell Signalling | 9282 | Mouse |
| RPA32-P-S4/S8 | WB | Bethyl laboratories | A300-245A | Rabbit |
| γH2AX | WB | Millipore | 07-164 | Rabbit |
| MEK2 | WB | BD | 610235 | Mouse |
| BrdU-FITC | FACS | BD | 556028 | Mouse |
| BrdU (CldU) | IF | Abcam | ab6326 | Rat |
| BrdU (IdU) | IF | BD | 347580 | Mouse |

**Supplementary Table 4.** Antibodies used in this study.

| Feature | Average size (bp) | % overlap | Source/definition |
|---|---|---|---|
| CGI | 655 | 10 | UCSC browser, (Gardiner-Garden and Frommer, 1987) |
| promoters | 2000 | 10 | RefSeq UCSC track, (Pruitt et al, 2004, 2014); +1.5 kb and - 0.5 kb from TSS |
| exons | 176.4 | 10 | RefSeq UCSC track, (Pruitt et al, 2004, 2014) |
| introns | 4619.8 | 10 | RefSeq UCSC track, (Pruitt et al, 2004, 2014) |
| TTS | 2000 | 10 | RefSeq UCSC track, (Pruitt et al, 2004, 2014); +/- 1kb from TTS |
| Intergenic regions | 92823.2 | 10 | RefSeq UCSC track (Pruitt et al, 2004, 2014) |
| G4 | 34 | 1bp | Home-made script; G4 were defined as GGGN[1,7]GGGN[1,7]GGGN[1,7]GGG, intersecting matches were merged |
| Early timing regions | $1.49 \times 10^{6}$ | 100 | Hiratani et al (2010) |
| Late timing regions | $2.21 \times 10^{6}$ | 100 | Hiratani et al (2010) |

**Supplementary Table 5. Genomic features intersected with origin datasets.** The overlap between an origin and a genomic feature was considered positive when the indicated minimum percentage of overlap was met.

| State | States as defined in Juan et al (2016) |
| --- | --- |
| Enhancers | 1+2+3+11+12+13+14 |
| Active promoters | 15+16+17 |
| Bivalent promoters | 18 |
| Transcriptional elongation | 4+5 |
| Insulators | 20 |
| PcG repressed | 19 |
| Heterochromatin | 6+7+8+10 |
| Low signal | 9 |

**Supplementary Table 6. Definition of chromatin states used in this study.**

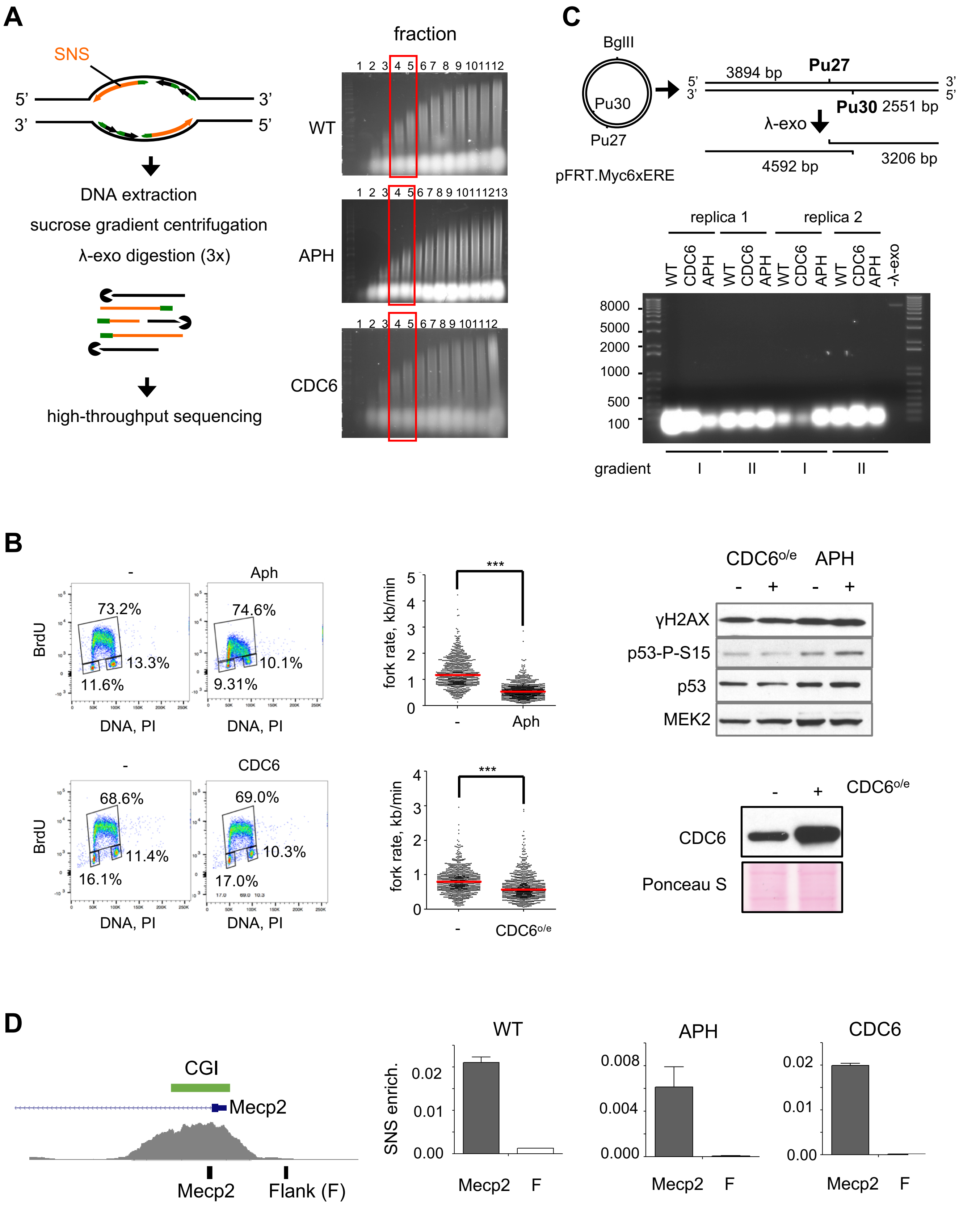
A
SNS
5'
3'
3'
5'
DNA extraction
sucrose gradient centrifugation
λ-exo digestion (3x)
high-throughput sequencing
fraction
WT
APH
CDC6
C
BglII
Pu30
Pu27
pFRT.Myc6xERE
3894 bp
Pu27
Pu30 2551 bp
λ-exo
4592 bp
3206 bp
replica 1
replica 2
WT CDC6 APH WT CDC6 APH WT CDC6 APH WT CDC6 APH -λ-exo
8000
5000
2000
1000
500
100
gradient
I
II
I
II
B
-
Aph
BrdU
73.2%
13.3%
11.6%
74.6%
10.1%
9.31%
DNA, PI
fork rate, kb/min
***
CDC6o/e
APH
γH2AX
p53-P-S15
p53
MEK2
CDC6
68.6%
11.4%
16.1%
69.0%
10.3%
17.0%
CDC6
Ponceau S
D
CGI
Mecp2
Flank (F)
WT
APH
SNS enrich.
Mecp2
F

### Supp Figure 2

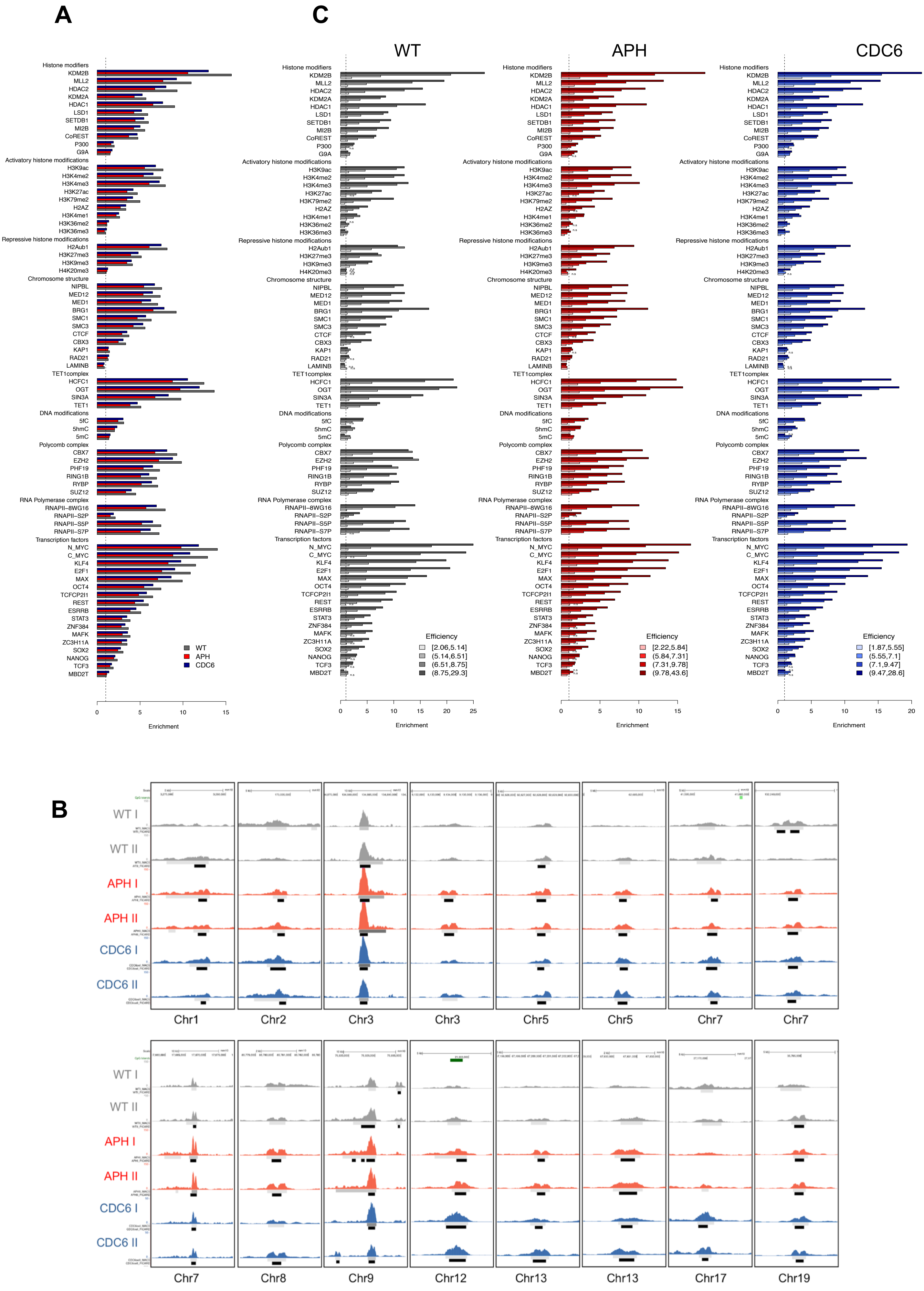

Supp Figure 3

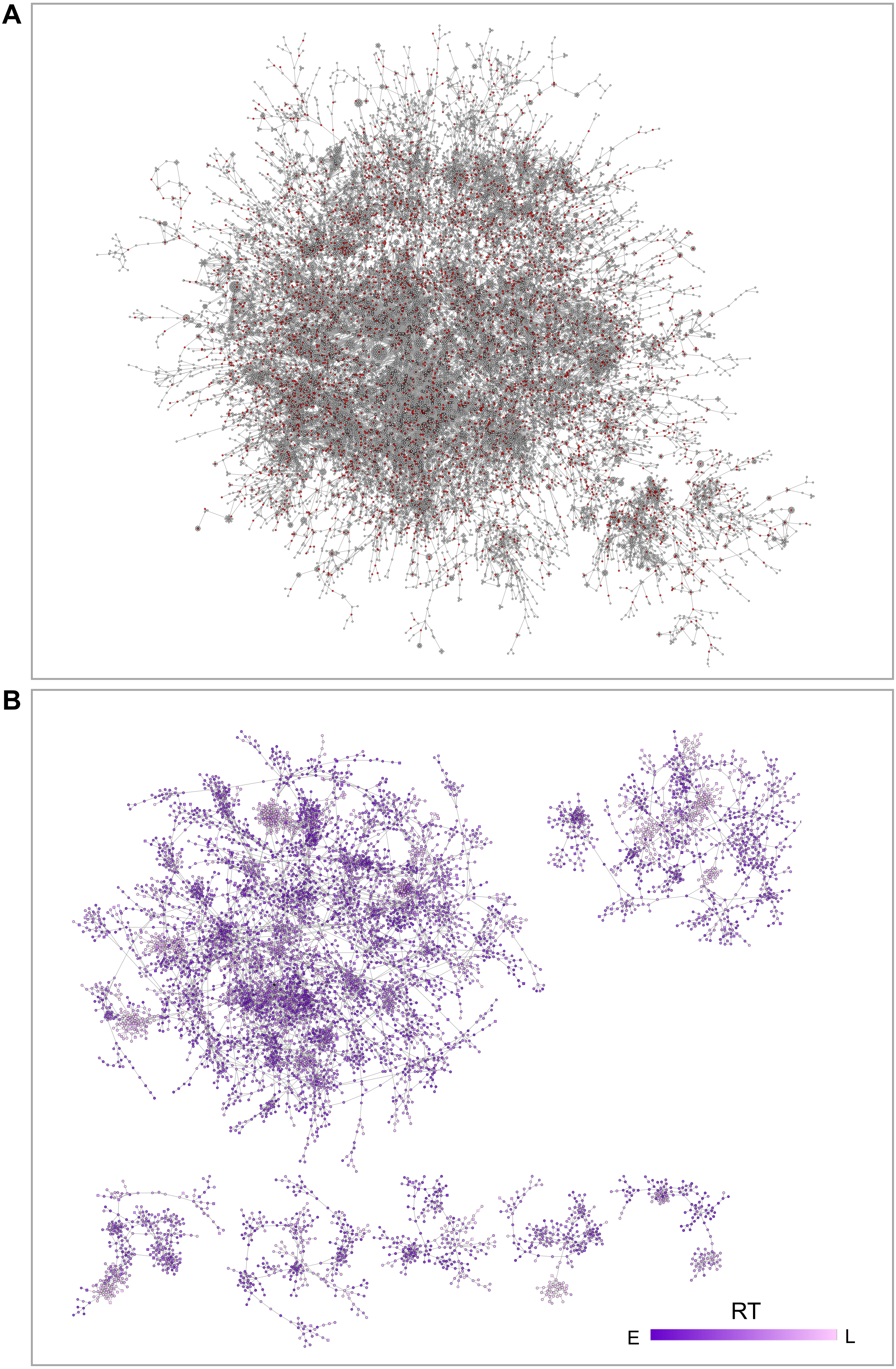

### Supp Figure 4

A

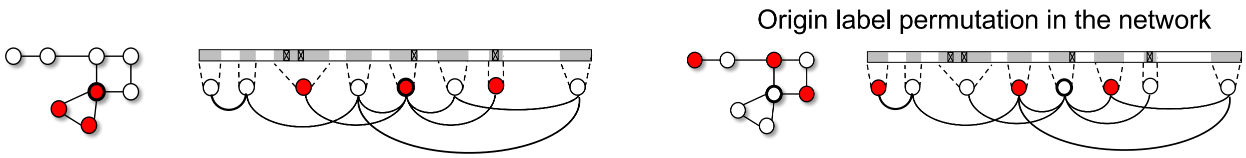


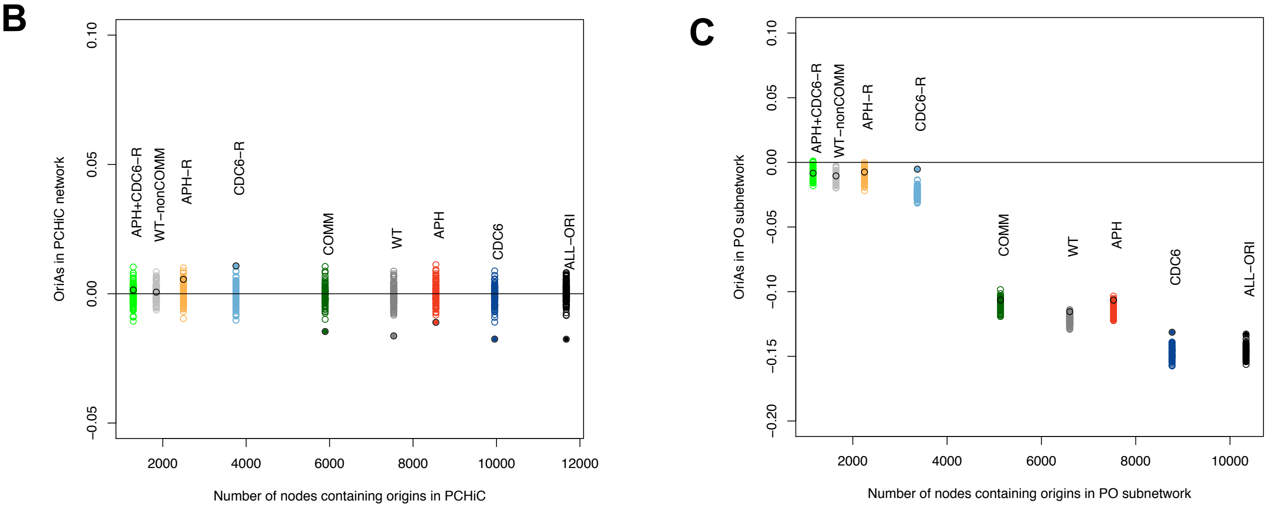


D

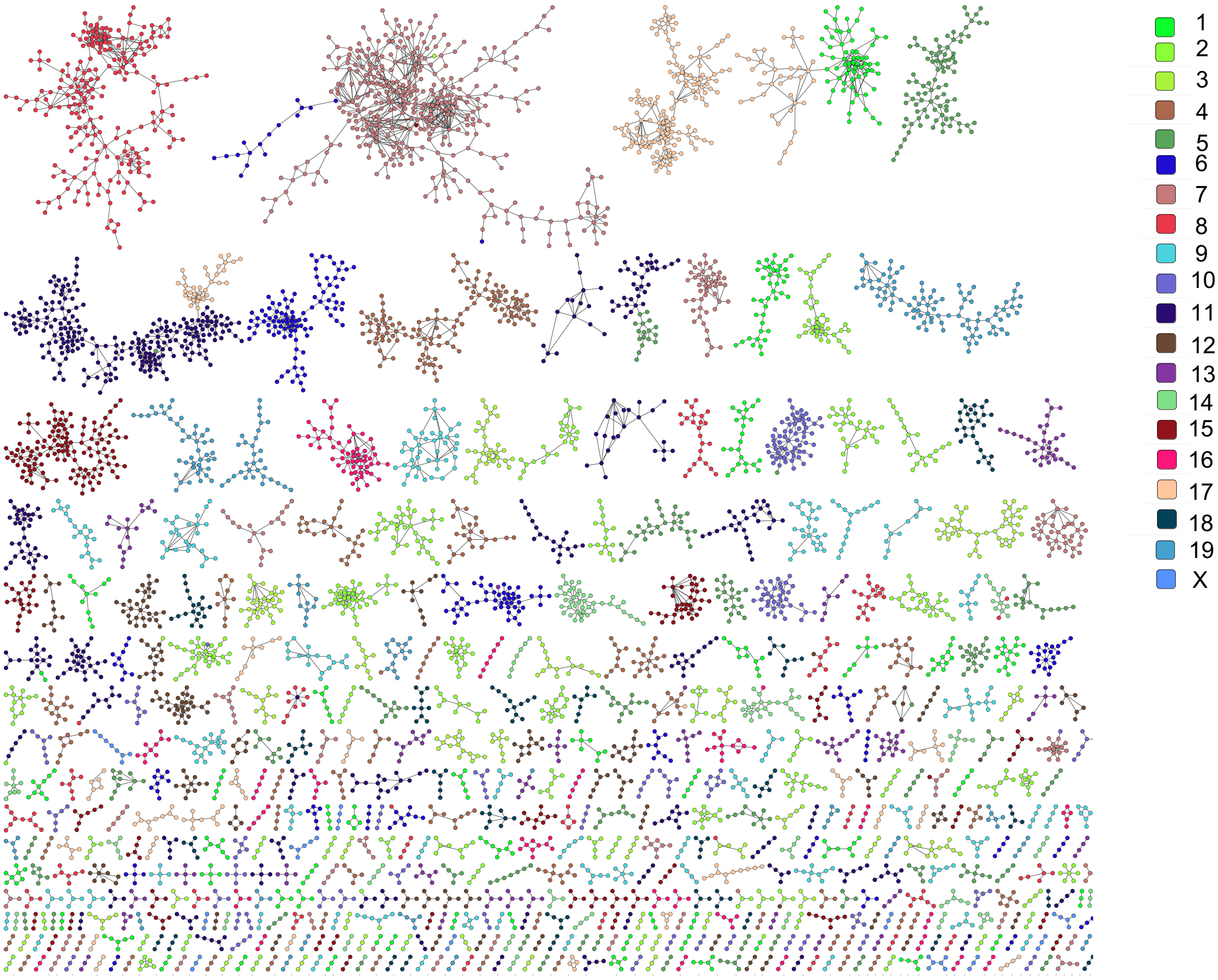

Supp Figure 5

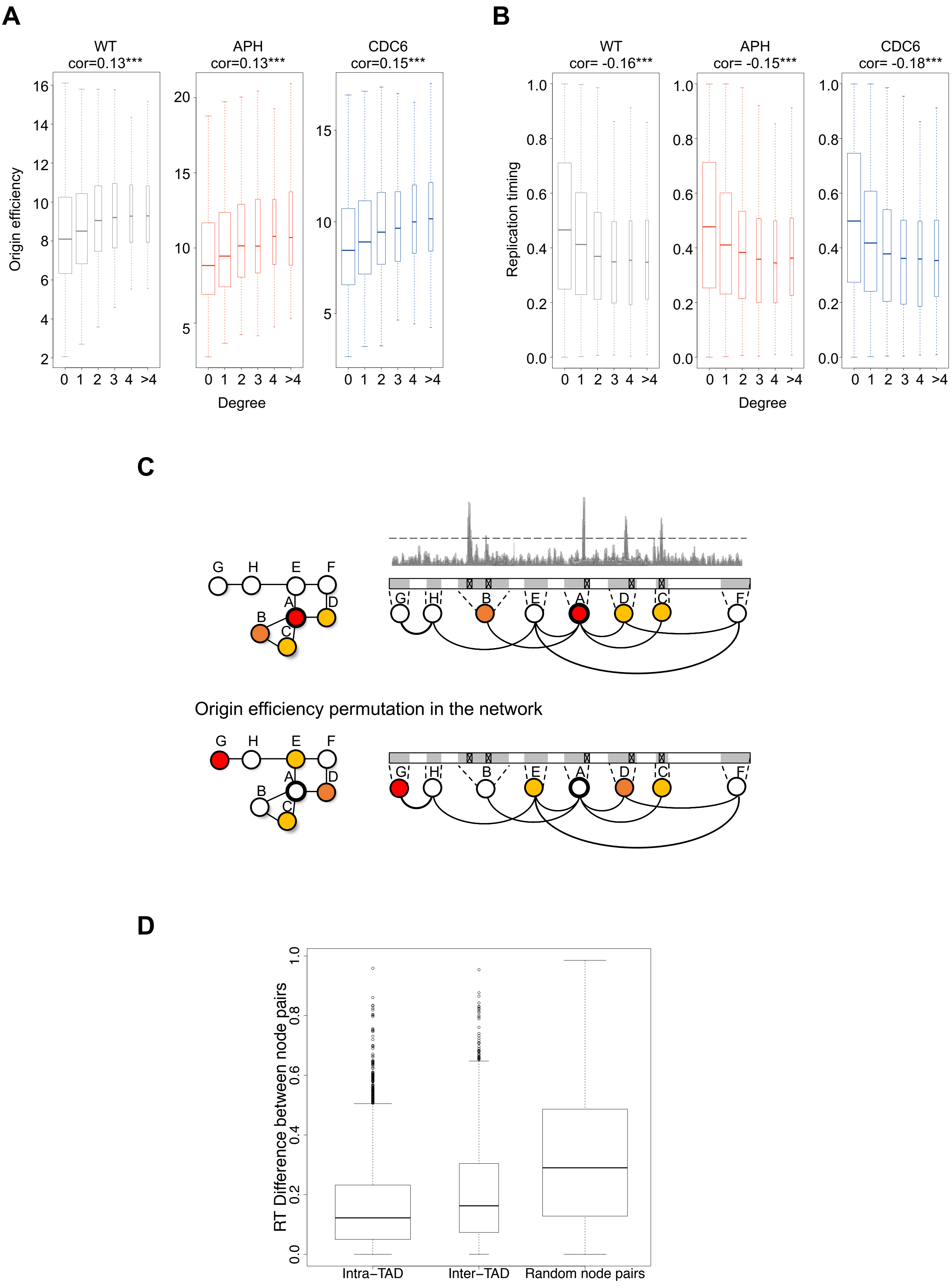

Supp Figure 6

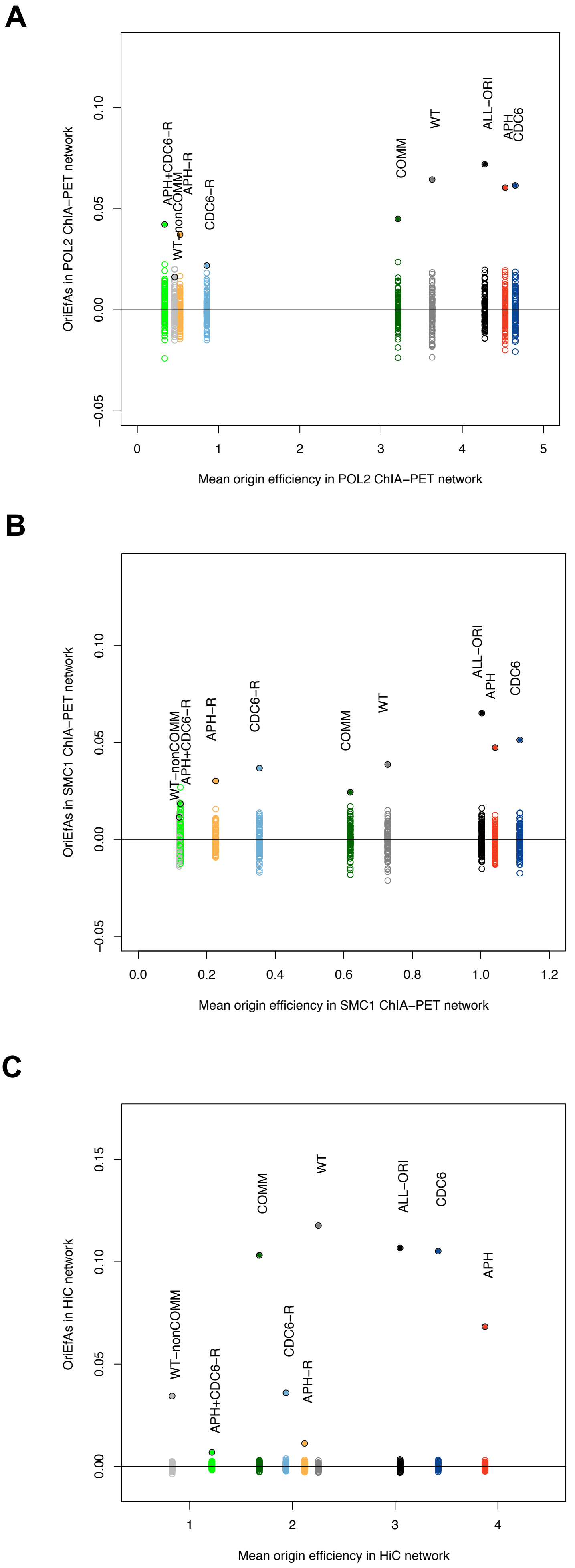
